# The microtubule regulator EFA-6 forms spatially restricted cortical foci dependent on its intrinsically disordered region and interactions with tubulins

**DOI:** 10.1101/2024.04.14.588158

**Authors:** Anjali Sandhu, Xiaohui Lyu, Xinghaoyun Wan, Xuefeng Meng, Ngang Heok Tang, Gilberto Gonzalez, Ishana N. Syed, Lizhen Chen, Yishi Jin, Andrew D. Chisholm

## Abstract

Microtubules (MTs) are dynamic components of the cytoskeleton and play essential roles in morphogenesis and maintenance of tissue and cell integrity. Despite recent advances in understanding MT ultrastructure, organization, and growth control, how cells regulate MT organization at the cell cortex remains poorly understood. The EFA-6/EFA6 proteins are recently identified membrane-associated proteins that inhibit cortical MT dynamics. Here, combining visualization of endogenously tagged *C. elegans* EFA-6 with genetic screening, we uncovered tubulin-dependent regulation of EFA-6 patterning. In the mature epidermal epithelium, EFA-6 forms punctate foci in specific regions of the apical cortex, dependent on its intrinsically disordered region (IDR). We further show the EFA-6 IDR is sufficient to form biomolecular condensates *in vitro*. In screens for mutants with altered GFP::EFA-6 localization, we identified a novel gain-of-function (gf) mutation in an α-tubulin *tba-1* that induces ectopic EFA-6 foci in multiple cell types. *tba-1(gf)* animals exhibit temperature-sensitive embryonic lethality, which is partially suppressed by *efa-6(lf)*, indicating the interaction between tubulins and EFA-6 is important for normal development. TBA-1(gf) shows reduced incorporation into filamentous MTs but has otherwise mild effects on cellular MT organization. The ability of TBA-1(gf) to trigger ectopic EFA-6 foci formation requires β-tubulin TBB-2 and the chaperon EVL-20/Arl2. The *tba-1(gf)-*induced EFA-6 foci display slower turnover, contain the MT-associated protein TAC-1/TACC, and require the EFA-6 MTED. Our results reveal a novel crosstalk between cellular tubulins and cortical MT regulators *in vivo*.

**Highlights:**

- The MT regulator EFA-6 forms spatially restricted punctate cortical foci
- The EFA-6 N-terminal intrinsically disordered region (IDR) is essential for the formation of cortical foci *in vivo* and is sufficient for droplet formation *in vitro*
- Tubulins regulate formation of EFA-6 foci via the EFA-6 MT elimination domain
- EFA-6 foci induced by altered tubulin heterodimer function display reduced turnover and recruit TAC-1/TACC

## INTRODUCTION

The cell cortex is a major site of regulation of microtubules (MTs) and their interaction with the actin-spectrin cytoskeleton. MTs interact with cortical proteins in various ways, including coupling to the dynein motor (Laan et al., 2012), CLASP proteins (Lawrence and Zanic, 2019), and spectraplakins (Nashchekin et al., 2016). Such dynamic interactions often stabilize MT ends at the cortex, counteracting the destabilizing effects of membrane contact. Other proteins inhibit cortical MT growth, including the Cdk1 kinase (Singh et al., 2021), the kinesin KIF18B (Moreci and Lechler, 2021), and the membrane-associated protein EFA-6/EFA6 (O’Rourke et al., 2010). EFA6 proteins contain a SEC7 domain that displays biochemical activity as an Arf6 GEF in mammalian cells (Franco et al., 1999; Sakagami, 2008) and associate with the plasma membrane via a PH domain. Membrane association has been shown to be important for EFA6’s Arf6 GEF activity (Kanamarlapudi, 2014); transmembrane proteins such as CD13 can also recruit EFA6 to punctate structures at the membrane (Ghosh et al., 2019). Functional studies in *C. elegans* and Drosophila have revealed that EFA6 restricts cortical MT growth in several types of cells, largely independent of its GEF activity (Chen et al., 2015; Chen et al., 2011; O’Rourke et al., 2010; Qu et al., 2019; Yogev et al., 2016). Key questions are how EFA6 inhibits cortical MT growth in different types of cells and how its MT-destabilizing activity is regulated in differentiated cells such as epithelia or neurons with extensive cortical MT arrays.

MTs are polymers of α/μ tubulin heterodimers. Decades of studies have elucidated intertwined mechanisms that ensure coordinated expression of tubulin heterodimers and their assembly into MTs with cell- and tissue-specific characteristics. Tubulin heterodimer levels homeostatically regulate tubulin expression at mRNA and protein levels (Gasic and Mitchison, 2019) and are controlled by the tubulin chaperone system (Al-Bassam, 2017). The soluble pool of tubulin heterodimers plays critical roles in autoregulation of tubulin synthesis, dynamics, and nucleation (Ohi et al., 2021). For example, centrosomes may act as microtubule organizing centers by locally concentrating tubulin heterodimers (Woodruff et al., 2017). Tubulin heterodimer binding also stimulates liquid-liquid phase separation and condensation of the centrosomal binding protein TPX2, leading to nucleation of noncentrosomal MTs (King and Petry, 2020). While some MT-regulating proteins may directly sense heterodimers, such as Stathmin 1 (Belmont and Mitchison, 1996; Ohi et al., 2021), it is unclear how MT organization within a cell is linked to local control of tubulin heterodimer levels.

Studies in *C. elegans* and Drosophila showed that the MT destabilizing function of EFA-6/EFA6 involves an N-terminal conserved motif (Chen et al., 2015; O’Rourke et al., 2010), named the MT elimination domain or MTED (Bu et al., 2021; Qu et al., 2019). The MTED is adjacent to a large N-terminal region predicted to be intrinsically disordered (Chen et al., 2015). The Drosophila EFA6 MTED can interact with unpolymerized tubulin in MT polymerization assays (Qu et al., 2019). Overexpression of Drosophila EFA6 N-terminal region containing both MTED and IDR can inhibit cortical MT growth in heterologous cell types. The MT inhibition function of EFA6 is also required for dendrite pruning *in vivo* (Bu et al., 2021). In *C. elegans* embryos, *efa-6(lf)* elevates cortical MT growth but does not cause major defects in cell division or differentiation. In neurons, *efa-6(lf)* mutants display excessive axon growth and increased MT dynamics (Chen et al., 2015; Chen et al., 2011), whereas overexpression of EFA-6 reduces axon growth and inhibits MT dynamics (Chen et al., 2015). In vertebrates, three EFA6 family members EFA6A/C/D are predominantly expressed in neurons (Eva et al., 2017; Franco et al., 1999), whereas EFA6B is mostly expressed in epithelia (Derrien et al., 2002) and has been implicated in maintaining epithelial morphology (Fayad et al., 2021). Like *C. elegans* and Drosophila, vertebrate EFA6 proteins contain large N-terminal IDRs. Less is known about whether vertebrate EFA6 family members regulate cortical MTs, and the mechanism of EFA6/MT regulation at the cortex has remained unclear.

Here we show that in *C. elegans* endogenously tagged EFA-6 localizes to the cellular cortex in many cell types, including embryonic blastomeres and multiple differentiated tissues. Strikingly, EFA-6 forms punctate cortical foci in the epidermal epithelium, dependent on its IDR. We show that the EFA-6 N-terminus can display condensation *in vivo* under stress and that a purified N-terminal IDR is sufficient to form biomolecular condensates *in vitro*. Through genetic screening, we find a gain of function (gf) mutation in an α-tubulin TBA-1 that triggers ectopic formation of stabilized EFA-6 foci. EFA-6 foci caused by TBA-1(gf) require TBB-2 β-tubulin and the tubulin chaperone EVL-20/Arl2 and are dependent on the EFA-6 MTED. Moreover, *tba-1(gf)-*induced EFA-6 foci recruit TAC-1/TACC, an EFA-6 binding partner and conserved component of pericentriolar material. Our findings reveal that the subcellular patterning of a cortical MT regulator is regulated by its interactions with specific tubulins *in vivo*.

## RESULTS

### EFA-6 localizes to the cellular cortex in multiple tissues and forms punctate foci in the apical epidermal cortex

To characterize endogenous EFA-6 expression, we generated knock-in (KI) strains expressing GFP or mScarlet (mSc) tagged EFA-6 at N-terminus using CRISPR-Cas9 engineering (Methods, Table S1). Previous studies suggested that N-terminal tagging does not interfere with EFA-6 function (Chen et al., 2015), and indeed GFP::EFA-6(*ju1658)* and mSc::EFA-6(*syb6998*) KI animals exhibited wild-type body shape, locomotion, growth rate, and fertility. *efa-6(lf)* mutants display overgrowth of touch receptor neurons (TRNs) (Chen et al., 2011), whereas *efa-6* KI animals displayed normal TRN axon growth and morphology (Figure S1A, B). GFP::EFA-6(*ju1658)* and mSc::EFA-6(*syb6998)* exhibited identical pattern, except that mSc::EFA-6 fluorescence quenched rapidly. For simplicity, we refer to either KI as EFA-6 in the text and state specific KI alleles in Figure legends.

Endogenously tagged EFA-6 proteins were widely expressed in multiple tissues, from one-cell embryos to adults. In early embryonic blastomeres (Figure 1A.vi, vii & S1C), EFA-6 localized to the cortex, consistent with prior studies of transgenically expressed EFA-6 (O’Rourke et al., 2007). In late larvae and adults, EFA-6 was detected in the epidermis, pharyngeal cells, body wall muscles, neuronal soma and processes, and syncytial germline (Figure 1A.i-v, S1C). In most cell types, EFA-6 showed an even distribution at the cellular cortex. In contrast, adult epidermal cells consistently displayed punctate cortical foci (‘foci’, 0.5-1.5 μm diameter) at the sublateral margins of lateral epidermal ridges; scattered smaller foci were also seen in the epidermis overlying muscle (Figure 1A.i-iii).

**Figure 1.**
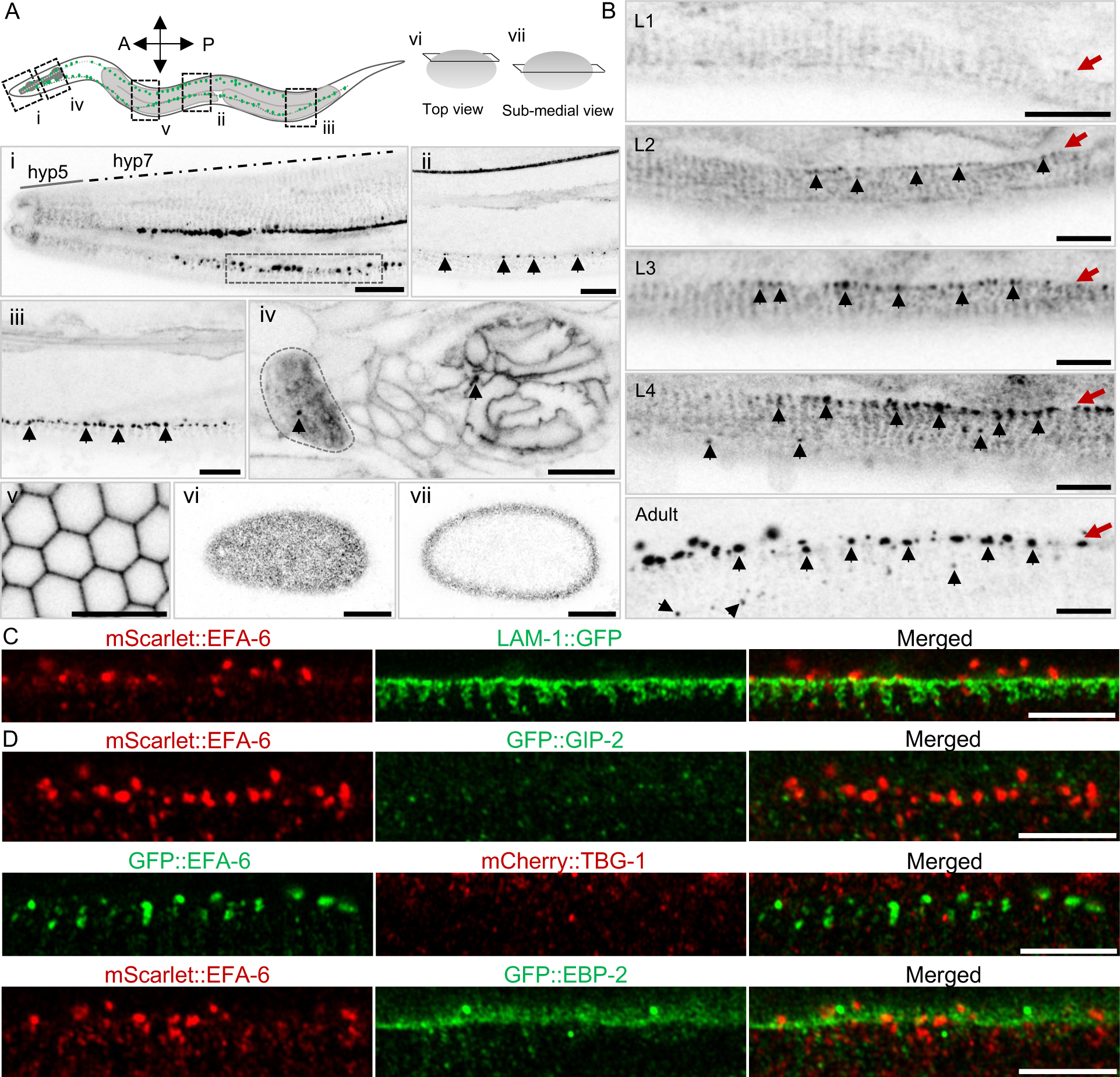
Endogenously expressed GFP::EFA-6 localizes to the cortex of multiple cell types and forms punctate foci along epidermal ridge margins. (A) Shown are single slice (0.5 µm thickness) confocal images of GFP::EFA-6(*ju1658*) knock-in in adults (i-v) and in single-cell stage embryos (vi, vii). Top cartoons show locations of regions corresponding to images in (i) anterior epidermis, (ii) mid-body epidermis, (iii) posterior epidermis, (iv) nerve ring/posterior bulb of the pharynx, and (v) mitotic germ line. (i-iii) GFP::EFA-6 displays circumferential bands in muscle adjacent to the epidermis and forms punctate foci (black arrows) along the dorsal and ventral margins of the lateral epidermal ridge. (i, ii) GFP::EFA-6 shows prominent localization along the length of the ALM process. (iv) GFP::EFA-6 shows a diffuse pattern in the nerve ring and localizes to the cortex of adjacent neuronal somas and pharyngeal muscle cells with rare punctate foci (black arrow). (v-vii) GFP::EFA-6 shows a diffuse pattern in developing oocytes and single-cell embryos. N = 5-15 animals per ROI. Scale = 10 µm. (B) Single slice confocal images of GFP::EFA-6 in the anterior lateral epidermis on the ventral side in larvae and adults, approximately corresponding to ROI (dashed box) in image A.i. GFP::EFA-6 foci (black arrows) begin to be detectable in the L3 stage and increase in number and intensity by the young adult stage. Red arrows indicate the epidermal ridge margins. N = 6, Scale = 5 µm. (C-D) Airyscan images show endogenously expressed mScarlet::EFA-6 foci along the lateral edge of the epidermal ridge localized near basal lamina component laminin β/LAM-1::GFP(*qyIs8*) (C), and such EFA-6 foci do not colocalize with MT minus-end protein γ-tubulin/mCherry::TBG-1(*ltSi62*), GFP::GIP-2(*lt19*), or with MT plus end binding protein EBP-2::GFP(*wow47*) (D). Images are single slices of 0.15 µm thickness, scale = 5 µm.

The larval and adult *C. elegans* epidermis is composed of four thickenings or ridges at lateral, dorsal, and ventral locations, connected by thinner sheets overlying body wall muscle quadrants (White et al., 1986); regions where the lateral ridges meet the muscle quadrants are here termed the sublateral ridge margins. EFA-6 foci at sublateral ridge margins were larger and more consistent than those in the epidermis overlying muscle, hence we analyzed the sublateral ridge margin foci in detail. EFA-6 foci at the sublateral ridge margins developed gradually through larval development, starting in the anterior hyp7 epidermis in the late L3 stage and extending along the length of the sublateral ridge margins by the first day of adulthood (Figure 1B). Formation of EFA-6 foci at the sublateral ridge margin formed within 20-25 µm of the anterior tip of the nose and extended posteriorly. EFA-6 foci were absent in the region of the epidermal cells hyp1-5 (Figure 1Ai). To clarify the subcellular localization of EFA-6 at the sublateral ridge margin, we examined colocalization with the basal lamina protein LAM-1 (Ziel et al., 2009). EFA-6 foci localized apical to LAM-1 (Figure 1C), suggesting EFA-6 foci reside in the apical cortex of the sublateral ridge margins.

The epidermis contains a complex MT network, with major MT bundles running longitudinally along lateral ridges as well as circumferentially (Chuang et al., 2016; Quintin et al., 2016; Wang et al., 2015). As axonal injury causes overexpressed EFA-6 to condense into cortical puncta that partially colocalize with MT minus end markers (Chen et al., 2015), we asked whether EFA-6 foci at the sublateral ridge margins could colocalize with MT end binding proteins. γ-tubulin TBG-1 and the γ-tubulin associated protein GIP-2 label MT minus ends in the apical epidermis (Wang et al., 2015); EFA-6 foci did not colocalize with TBG-1 or GIP-2 (Figure 1D). The MT plus end binding protein EBP-2 accumulates in longitudinal bands at the sublateral ridge margin (Castiglioni et al., 2020); EFA-6 foci were occasionally partially overlapping but largely did not colocalize with EBP-2 (Figure 1D). These data suggest that in the epidermis, EFA-6 cortical foci generally do not colocalize with MT ends.

### The IDR is necessary for formation of EFA-6 foci *in vivo* and sufficient for condensate formation *in vitro*

We next addressed which domains of EFA-6 were necessary for the formation of cortical foci at the sublateral ridge ridge margin using genome editing to delete or mutate coding sequences for specific domains (Methods, Table S2-3). The N-terminal region of EFA-6 is essential for its function in embryonic cytokinesis (O’Rourke et al., 2010) and neuronal axon regeneration (Chen et al., 2015). The EFA-6 MTED (aa 25-42) is predicted in Alphafold to form a beta-sheet hairpin loop (Figure S2A). EFA-6 N-terminal region also contains a large predicted intrinsically disordered region (IDR) that we divide into IDR1 and IDR2 for simplicity (Figure S2A, C). We primarily used GFP::EFA-6 for genome editing because of its stable fluorescence; key results were confirmed with mSc::EFA-6. Deletion of exons encoding the EFA-6 C-terminal domains resulted in transcript instability and low protein expression. Nonetheless, deletion of the PH and C-terminal coiled-coil domain (ΔPH+CC, *ju1833*) caused the remaining GFP::EFA-6 to be cytosolic (Figure S3A), consistent with the PH domain conferring cortical localization. We made multiple in-frame deletion alleles affecting the N-terminus of EFA-6 and also edited the highly conserved amino acid residues in MTED, Ser33 and Gly34 (Figure S2B, C). The overall pattern and intensity of GFP::EFA-6 in these edited alleles resembled those of unedited GFP::EFA-6 in most tissues (Figure S2D-F). Deletion of the N-terminal 176 amino acids, which include MTED and IDR1 (ΔMTED+IDR1, *ju1903*), eliminated GFP::EFA-6 foci at the epidermal ridge (Figure 2B, H). However, mutating S33 (*ju1974*) or S33 and G34 together in MTED (*ju2011*) did not alter GFP::EFA-6 foci formation (Figure 2C-D, H). Deleting the entire IDR (ΔIDR1+IDR2, *ju1990*) eliminated GFP::EFA-6 foci at the epidermal ridge, similar to EFA-6(ΔMTED+IDR1), while deletion of IDR1(*ju2026*), but not IDR2(*ju1943*), reduced the number and intensity of GFP::EFA-6 foci at the sublateral ridge margin (Figure 2A, E-I). These data suggest the N-terminal segment of the IDR, particularly IDR1, is necessary for EFA-6 cortical foci formation in the epidermis.

**Figure 2:**
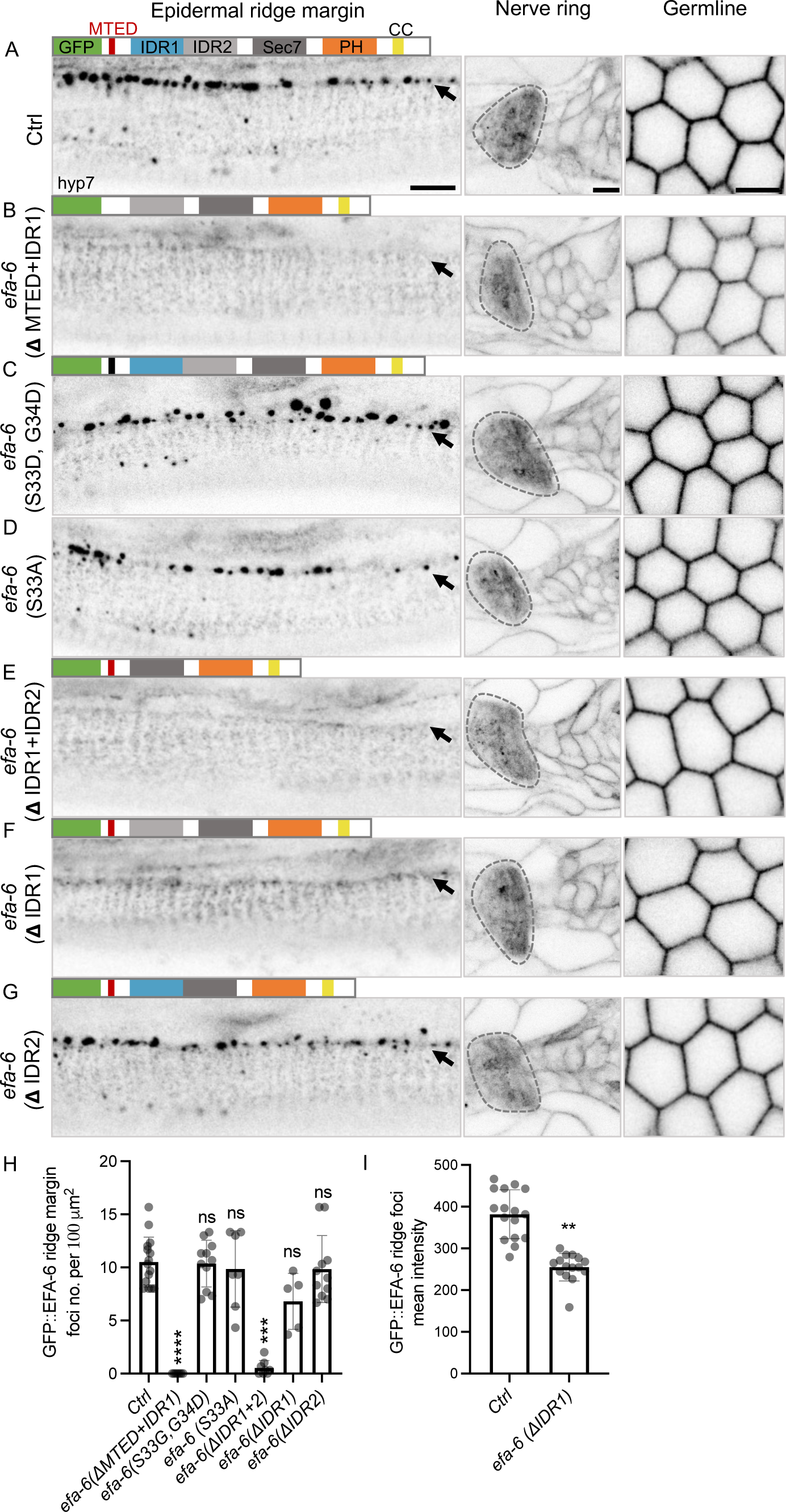
GFP::EFA-6 cortical foci formation requires its intrinsically disordered region (IDR) (A-G) Shown are confocal single slice (0.5 µm thickness) images of full-length and CRISPR-edited GFP::EFA-6 mutants in the anterior epidermal ridge margin (hyp7 cell, margin indicated by black arrows), nerve ring, and germline, with corresponding GFP::EFA-6 protein illustration above the image and allele marked on the left side. Scale = 5 µm. (H) Quantitation of GFP::EFA-6 cortical foci density along the ventral sublateral margin of the epidermal ridge. (I) Quantitation of mean fluorescence intensity of GFP::EFA-6 cortical foci along the epidermal ridge margin. (H-I) N = 5-15 animals; Error bar, SEM; Statistics, Kruskal-Wallis test is used in H and unpaired t-test in I, ns (not significant), ** P ≤ 0.01, *** P ≤ 0.001, **** P ≤ 0.0001.

Many proteins containing intrinsically disordered regions undergo phase separation under stress conditions (Uversky, 2017). We assessed if EFA-6 localization was affected by mild heat stress (2 h at 34°C; Methods), a condition that induces phase separation of other proteins in *C. elegans* (Andrusiak et al., 2019). We subjected both full-length GFP::EFA-6 and C-terminal truncated GFP::EFA-6(ΔPH+CC, *ju1833*) animals to heat stress. We observed that heat stress consistently caused GFP::EFA-6(ΔPH+CC) to form cytosolic foci, while the pattern of full-length GFP::EFA-6 remained the same (Figure S3A). The heat-induced GFP::EFA-6(ΔPH+CC) foci disappeared gradually over 8 h recovery at room temperature (Figure S3B). These observations are consistent with the N-terminus of EFA-6 being capable of phase separation under heat stress. We then tested whether the EFA-6 N-terminal region can form condensates using an *in vitro* droplet assay (Chen et al., 2023) (Methods). We expressed and purified GFP or mCherry fusions to the EFA-6 N-terminal 150 aa (EFA-6N), corresponding approximately to the coding sequence deleted in EFA-6(ΔMTED+IDR1). Purified GFP::EFA-6N or mCh::EFA-6N protein formed spherical droplets *in vitro* whose size was dependent on concentration (Figure S3C-D). Treatment with 1,6-hexanediol, which disrupts multivalent hydrophobic interactions in phase-separated condensates (Kato and McKnight, 2018), was able to dissolve EFA-6N droplets. Together, these data show that the EFA-6 IDR is capable of phase separation to form biomolecular condensates *in vitro*. We infer that under normal physiological conditions, the cortical localization of full-length EFA-6 generally restricts its propensity to form localized condensates except at the epidermal ridge margin. Henceforth we refer to the *in vivo* GFP::EFA-6 cortical foci as condensates.

### A gain-of-function mutation in α-tubulin induces ectopic EFA-6 condensates

To understand how endogenous EFA-6 localization is regulated, we conducted a forward genetic screen for altered GFP::EFA-6 expression (Methods). We isolated a mutant, *ju1761*, showing widespread ectopic GFP::EFA-6 condensates in many tissues throughout embryonic development and in adults (Figure 3A). We mapped this mutation to a single nucleotide alteration in the α-tubulin *tba-1,* changing Arg 241 to Cys (R241C). We observed identical effects on GFP::EFA-6 or mSc::EFA-6 when we made the same nucleotide change by genome editing (Methods and Tables S1-3), hereby referred to as *tba-1(R241C)*. *tba-1(R241C)* did not strongly affect the density of EFA-6 condensates at the epidermal ridge margin; however, additional dispersed condensates were observed in the epidermis, germ cells, and neurons (Figure 3). Notably, *tba-1(R241C)* mutants displayed abundant ectopic EFA-6 condensates in the head tip containing hyp5 and more anterior epidermal cells (Figure 3A). As proteins with intrinsically disordered sequences are prone to form condensates when overexpressed (Chakrabortee et al., 2016), we also examined total EFA-6 protein levels by western blotting and found no detectable increase in *tba-1(gf)* compared to controls (Figure S4A).

**Figure 3.**
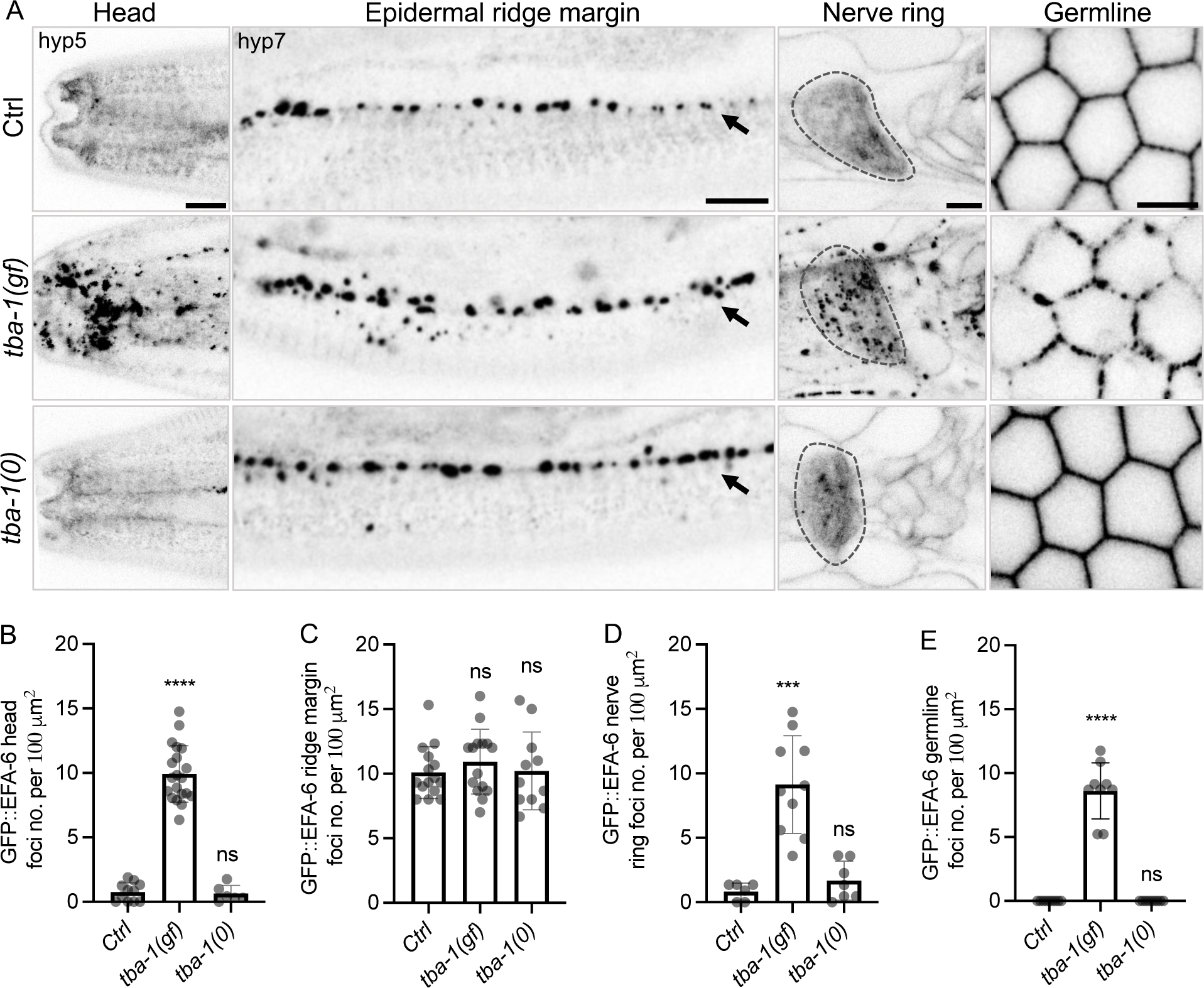
A gain of function mutation in α-tubulin TBA-1 induces ectopic GFP::EFA-6 condensates formation. (A) Confocal single slice images of GFP::EFA-6 in the head epidermis (hyp5), anterior epidermal ridge margin (hyp7, arrows), nerve ring, and syncytial germline in control, *tba-1*(gf, R241C) and *tba-1(0)* null mutants (day 1 adults); scale = 5 µm. (B-E) Quantitation of GFP::EFA-6 condensates density in the head (B), lateral epidermal ridge margin (C), nerve ring (D), and germline (E). Each dot represents EFA-6 condensates density in one ROI, normalized to 100 µm^2^ (mean and SEM); N = 6-9 animals per genotype; Kruskal-Wallis test, ns (not significant), ** P ≤ 0.01, *** P ≤ 0.001, **** P ≤ 0.0001.

Unlike *tba-1(R241C)*, *tba-1(0)* null mutants showed normal EFA-6 localization in all cell types examined, and the pattern and density of EFA-6 condensates at the epidermal ridge margin was comparable to control (Figure 3). We also knocked down *tba-1* by RNA interference (RNAi) in *tba-1(R241C)* mutants and observed reduced EFA-6 condensate formation (Figure S4B). *tba-1(R241C)* behaved as a dominant mutation, as we observed EFA-6 condensates in heterozygous animals *(tba-1(R241C)/+)* resembling that of *tba-1(R241C)* homozygotes (Figure S4C). These data indicate that the R241C missense change causes a gain of function in TBA-1; we refer to it as below *tba-1(gf)*. Under normal culture temperature (20°C), *tba-1(gf)* mutants displayed grossly normal body morphology, movement, and growth rate. However, when cultured at 25°C, *tba-1(gf)* exhibited 100% embryonic lethality (Table 1). As EFA-6 is involved in embryonic cortical MT organization (O’Rourke et al., 2010), we assessed whether *efa-6* interacted with *tba-1(gf)* in embryonic development. We found that both *efa-6(0)* null and *efa-6*(μIDR1+IDR2) mutations significantly rescued temperature-sensitive embryonic lethality of *tba-1(gf)*, while *efa-6*(μMTED+IDR1) mutation exhibited similar, but weaker, effects (Table 1). These genetic results suggest that *tba-1(gf)* and *efa-6* function antagonistically in MT organization, dependent on the EFA-6 IDR.

**Table 1.**
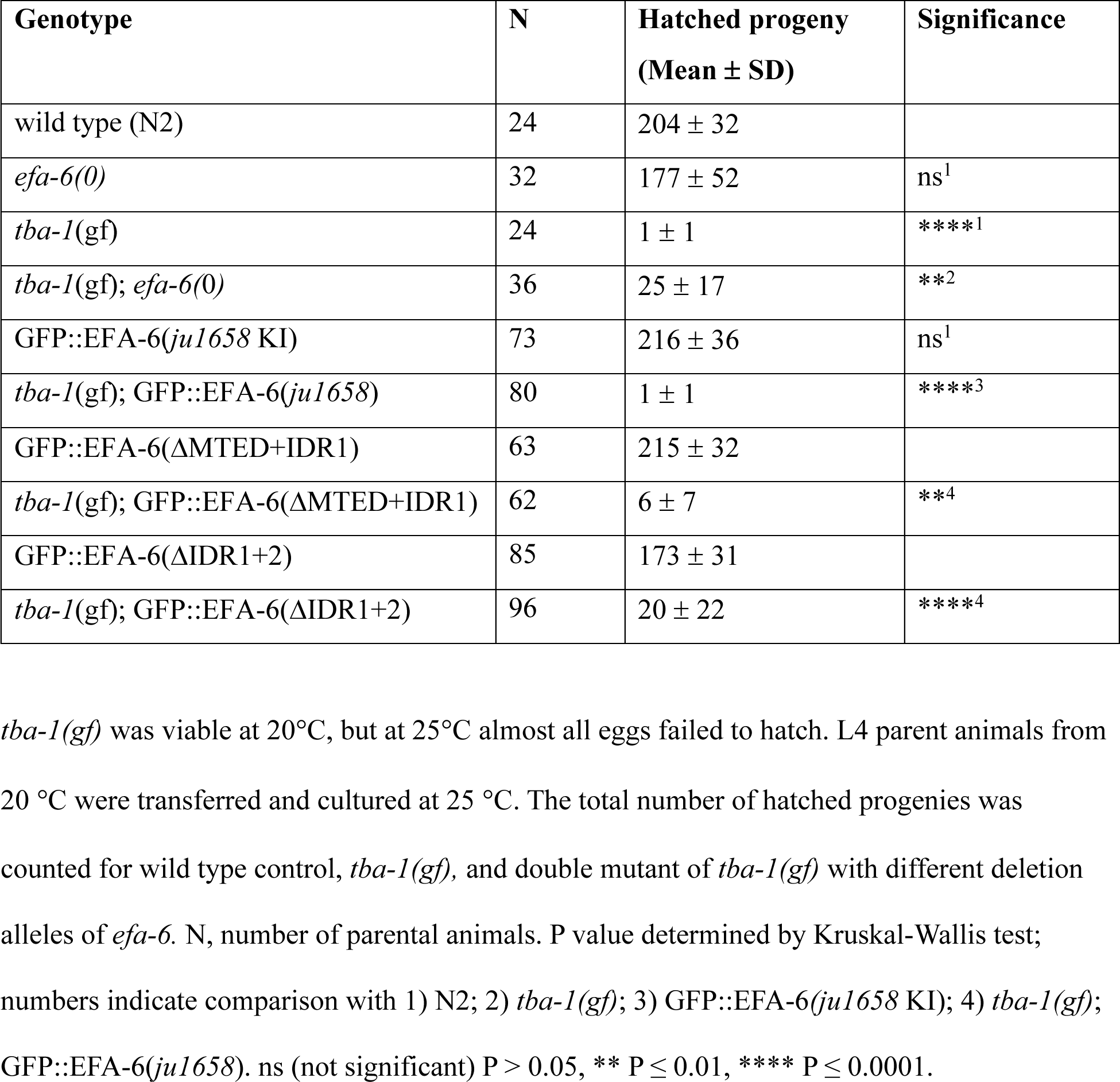
*efa-6* loss of function suppresses embryonic lethality of *tba-1(gf)*

### α-tubulin R241C induced EFA-6 condensate formation depends on **β-**tubulin TBB-2 and the chaperone Arf-like GTPase EVL-20

R241 is invariant in all α-tubulins and is located in the H7 helix close to the inter-dimer interface (Chaaban et al., 2018) (Figure S4D). Based on structural studies of tubulins (Alushin et al., 2014), TBA-1 R241 is predicted to be largely buried, with its side chains contacting multiple other amino acid residues. We next asked if the effect of TBA-1(R241C) is due to the absence of arginine or the presence of cysteine. We generated multiple edited *tba-1* alleles, changing R241 to alanine (neutral), lysine (positive), or glutamate (negative). We found all these mutations caused similar ectopic EFA-6 condensate formation, with varying degrees of embryonic and maternal-effect lethality (Table 2). Both TBA-1(R241K) and TBA-1(R241E) showed non-conditional maternal-effect lethality, whereas TBA-1(R241A) displayed temperature-sensitive embryonic lethality, resembling TBA-1(R241C) (Table 2). In contrast, editing of L240E in the H7 helix of TBA-1 did not cause ectopic GFP::EFA-6 condensates or embryonic lethality. These data indicate a crucial role of R241 in α-tubulin function. Another α-tubulin TBA-2 is widely coexpressed with TBA-1. We made a R241C mutation in TBA-2 and observed that TBA-2(R241C) showed ectopic EFA-6 condensate formation indistinguishable from that in TBA-1(gf). Moreover, TBA-2(R241C) exhibited non-conditional maternal-effect embryonic lethality (Table 2), possibly reflecting that TBA-2 is more abundant than TBA-1 (Nishida et al., 2021). These data identify the R241 residue in α-tubulins as critical in repressing EFA-6 condensate formation in both mitotic and differentiated cells.

**Table 2.** Phenotypes of *tba-1* and *tba-2* missense mutants.

| Amino acid change | Ectopic GFP::EFA-6 condensation | Gross Phenotypes | % eggs hatched at 20°C | % eggs hatched at 25°C |
| --- | --- | --- | --- | --- |
| *tba-1* R241C | 100% | Superficially WT at 20 °C, mild Unc at 25 °C | ~80 | 0 |
| *tba-1* R241A | 100% | Superficially WT at 20°C, mild Unc at 25°C | ~100 | 0 |
| *tba-1* R241K | 100% | Small and lay dead eggs at 20 °C, mild Unc at 25°C | 0 | 0 |
| *tba-1* R241E | 100% | Small and lay dead eggs at 20°C, mild Unc at 25°C | 0 | 0 |
| *tba-1* L240E | 0% | Superficially WT at 20 °C and 25 °C | 100 | 100 |
| *tba-2* R241C | 100% | Small and lay dead eggs at 20°C, mild Unc at 25°C | 0 | 0 |

L4 parental animals were cultured at 20°C or 25°C and the number of hatched progeny counted after 24 h. N = 4 parents per genotype per trial, 3 independent trials.

β-tubulins are obligate functional partners of α-tubulins. Most types of cells express several tubulin isotypes, with each α-tubulin forming partners with multiple β-tubulins (Janke and Magiera, 2020). Moreover, a complex set of tubulin-specific chaperones regulates tubulin heterodimer formation and incorporation into polymers (Al-Bassam, 2017). In *C. elegans,* TBB-1 and TBB-2 are two major β-tubulins widely coexpressed with TBA-1 and can form heterodimers with both TBA-1 and TBA-2 (Chaaban et al., 2018; Honda et al., 2017). To address if TBA-1(gf) requires a specific β-tubulin to induce ectopic EFA-6 condensation, we examined how loss of function mutation in *tbb-1* and *tbb-2* affected *tba-1(gf)*. We found that while neither *tbb-1(0)* or *tbb-2(0)* affected EFA-6 cortical condensates at the epidermal ridge margin, *tbb-2(0)*, but not *tbb-1(0)*, specifically alleviated the ectopic EFA-6 condensates in *tba-1(gf)* mutants (Figure 4A-F, I-L). *evl-20* is orthologous to the tubulin dimerization chaperone Arl2 (Antoshechkin and Han, 2002). *evl-20(0)* mutants showed grossly normal EFA-6 condensates at the epidermal ridge margins and also significantly suppressed the ectopic EFA-6 condensates in *tba-1(gf)* mutants (Figure 4A, G, H, I-L). These findings suggest the ability of TBA-1(gf) to induce ectopic EFA-6 condensates depends on tubulin heterodimer formation, specifically with TBB-2.

**Figure 4.**
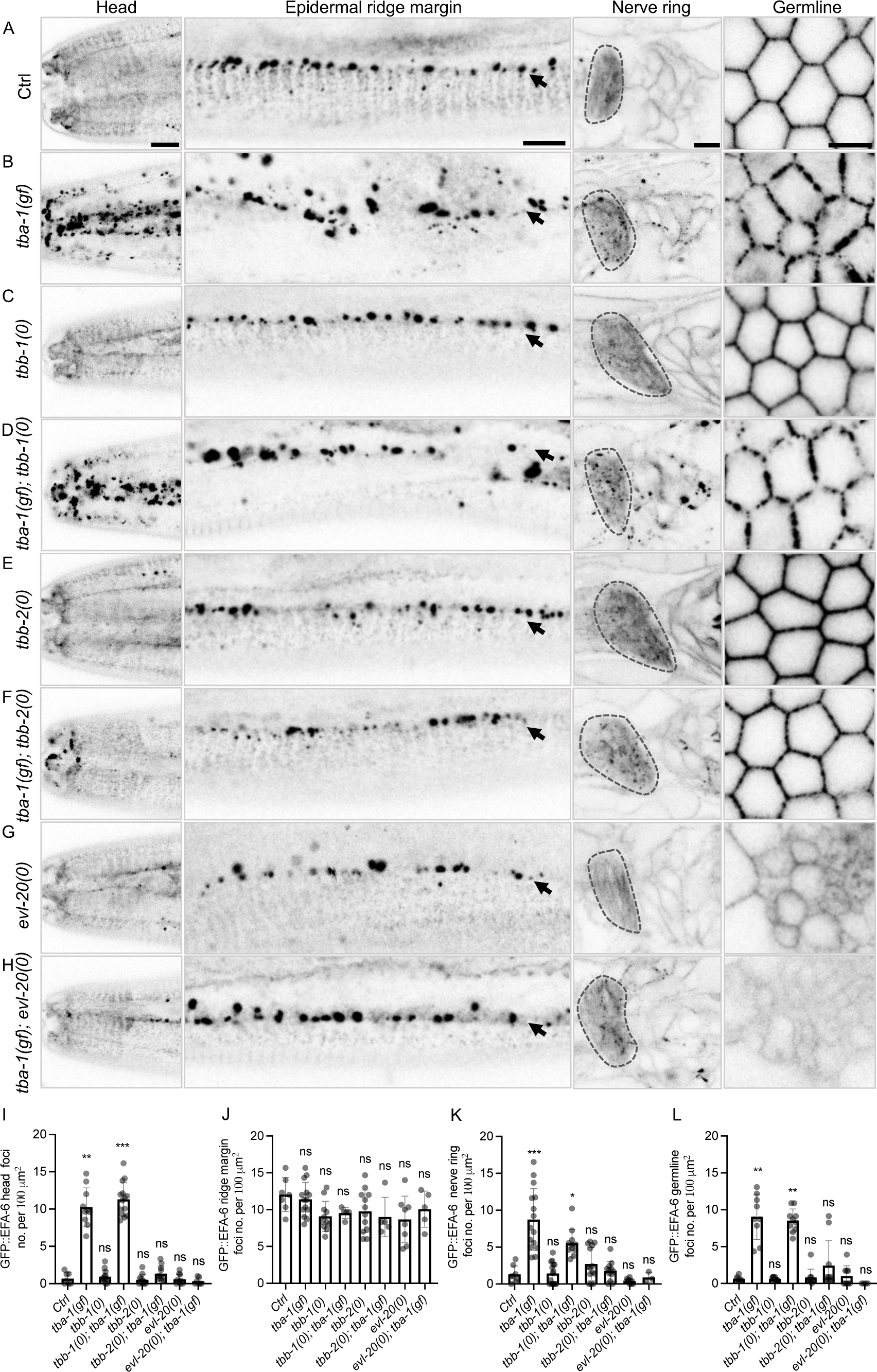
*tba-1(gf)* induced GFP::EFA-6 ectopic condensates require TBB-2 β-tubulin and the tubulin chaperone EVL-20. (A-H) Confocal single slice images of GFP::EFA-6 in control and mutants with genotypes indicated on the left. Black arrowheads point to the lateral epidermal ridge margin; Scale = 5 µm. (I-L) Quantitation of GFP::EFA-6 condensate (foci) density in the head (I), lateral epidermal ridge margin (J), nerve ring (K), and germline (L). Each dot represents a single animal; N = 5-10 animals per genotype. Data represent mean and SEM; P value determined by Kruskal-Wallis test, ns (not significant) P > 0.05, * P ≤ 0.05, ** P ≤ 0.01, *** P ≤ 0.001, **** P ≤ 0.0001.

### TBA-1(R241C) displays reduced incorporation into polymerized microtubules

We next asked how *tba-1(gf)* might affect overall MT organization. We first examined the expression pattern and intensity of several MT-binding proteins in the epidermis. The MT shaft binding protein MAPH-1.1(KI) displayed a filamentous pattern, labelling MTs along the epidermal ridge as well as in the circumferential MT bundles. In *tba-1(gf)* mutants, the overall pattern and the number of MAPH-1.1-labelled MT bundles were comparable to that in control, although the total intensity of MAPH-1.1 was weaker (Figure 5A-C). We also visualized MT organization by GFP::TBB-1(KI) and GFP::TBB-2(KI) and observed no obvious effects in *tba-1(gf)* compared to wild type (Figure S5A, B). These data suggest that *tba-1(gf)* does not cause major disruption of overall MT architecture, possibly due to functional compensation by other tubulins.

**Figure 5.**
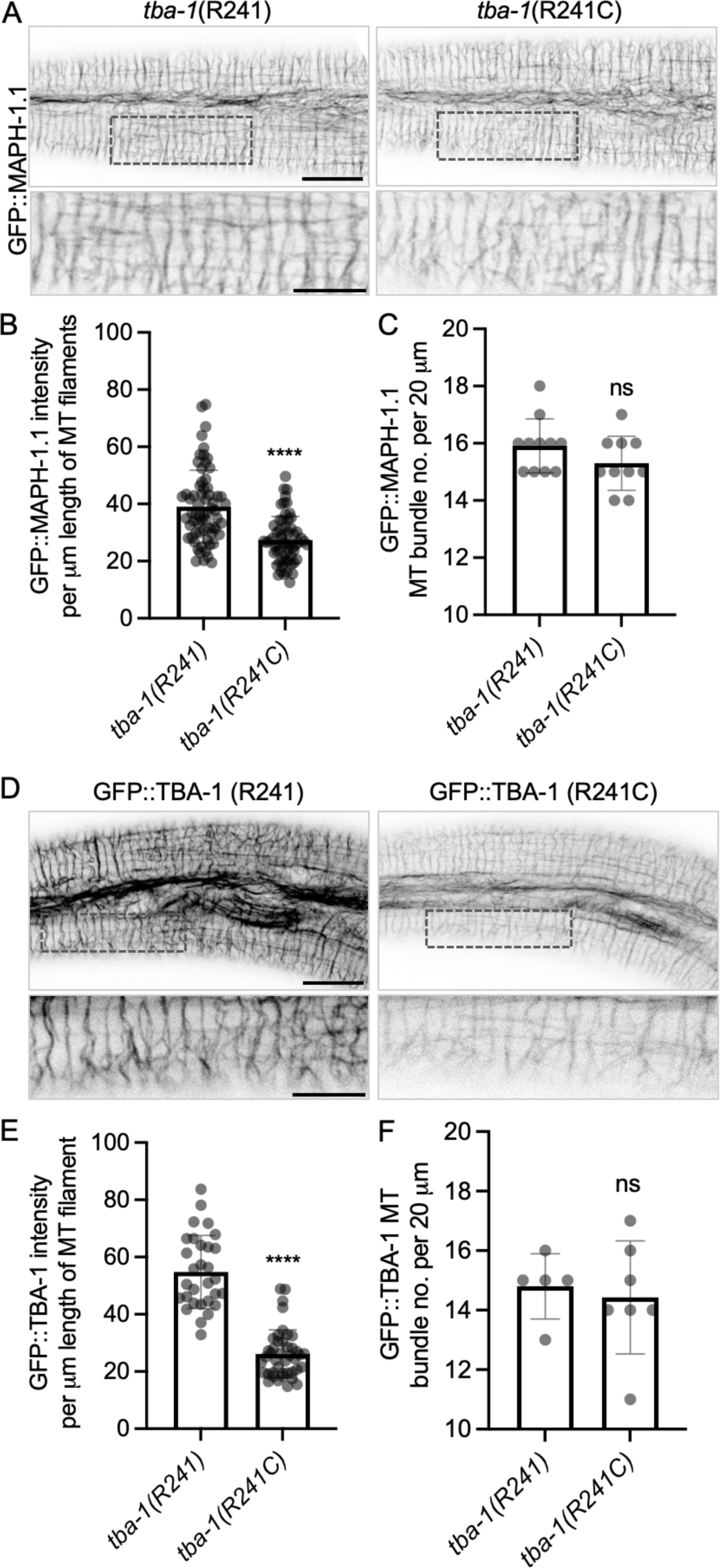
*tba-1(gf)* shows reduced incorporation into polymerized MTs. (A) Shown are confocal single slice images of MT shaft marker GFP::MAPH-1.1(*mib12*) in control *tba-1(R241)* and *tba-1(R241C, gf)* animals (1-day adult). Dashed boxes correspond to the enlarged view below. Scale = 10 µm for top images and 5 µm for enlarged images. (B, C) Quantitation of GFP::MAPH-1.1-labeled circumferential MT bundles and number of MT bundles in a 20 µm region in the epidermis (N = 10). (D) Confocal single slice images show reduced incorporation of GFP::TBA-1(R241C, gf) into epidermal MT bundles, compared GFP::TBA-1(R241, +); dashed boxes correspond to enlarged view below. (E, F) Quantitation of GFP::TBA-1(R241, +) and GFP::TBA-1(R241C, gf) labeled circumferential MT bundles and number of MT bundles in a 20 µm region in the epidermis of day 1 adults (N = 5-8). Error bar in graphs represents SEM; P value determined by Mann-Whitney test, ns (not significant) P > 0.05, **** P ≤ 0.0001.

We further asked how TBA-1(gf) affects its incorporation into MTs. Endogenously tagged GFP::TBA-1(KI) displays a filamentous pattern, similar to GFP::MAPH-1.1, in the epidermis (Figure 5D) and other tissues throughout development (Honda et al., 2017). We edited the R241C mutation in the GFP::TBA-1(KI) and found the overall fluorescence intensity of GFP::TBA-1(gf) in epidermal MT bundles was significantly lower than that of GFP::TBA-1(+), although the number of MT bundles remained similar (Figure 5D-F). Unlike untagged *tba-1(gf)*, however, GFP::TBA-1(gf) animals did not display ectopic EFA-6 condensates or embryonic lethality at 25 °C, consistent with evidence that GFP tagging impairs TBA-1 function, as found for other *C. elegans* tubulins (Honda et al., 2017). As an independent test of the ability of TBA-1(gf) to incorporate into MTs, we took advantage of the fact that *tba-1(gf)* is genetically dominant (Figure S4C). We compared GFP::TBA-1 intensity in *GFP::TBA-1*/*tba-1(gf)* animals to *GFP::TBA-1*/*tba-1(+)* and found that GFP::TBA-1 intensity was higher when in *trans* to *tba-1(gf)* than when in trans to *tba-1(+)* (Figure S5C). These comparisons suggest TBA-1(gf) is less efficiently incorporated into MTs compared to TBA-1(+).

### The EFA-6 MTED is required for induction of ectopic EFA-6 condensates by TBA-1(R241C)

Next, we addressed which domains of EFA-6 are required for the formation of ectopic condensates in *tba-1(gf)* mutants. We generated double mutants of *tba-1(gf)* with several mutant versions of GFP::EFA-6 (Figure S2, Table S1). GFP::EFA-6(ΔMTED+IDR1) did not undergo condensation in *tba-1(gf)* mutants (Figure 6C). However, deletion of the entire IDR (ΔIDR1+IDR2), which abolished EFA-6 condensate formation in the wild type (Figure 2), reduced but did not eliminate ectopic EFA-6 condensation in *tba-1(gf)* mutants (Figure 6C, H). In *tba-1(gf)* animals, GFP::EFA-6 lacking either IDR1 or IDR2 also formed ectopic condensates indistinguishable from *tba-1(gf)* mutants (Figure 6D, E), suggesting *tba-1(gf)* can induce ectopic EFA-6 condensation partially independent of the IDR. We then tested the MTED mutant GFP::EFA-6(S33D, G34D) and found no ectopic condensates in *tba-1(gf)* mutants (Figure 6F). This suggests that although the MTED is not required for formation of normal EFA-6 condensates at the epidermal ridge margin, it is necessary for *tba-1(gf)*-induced ectopic EFA-6 condensate formation.

**Figure 6.**
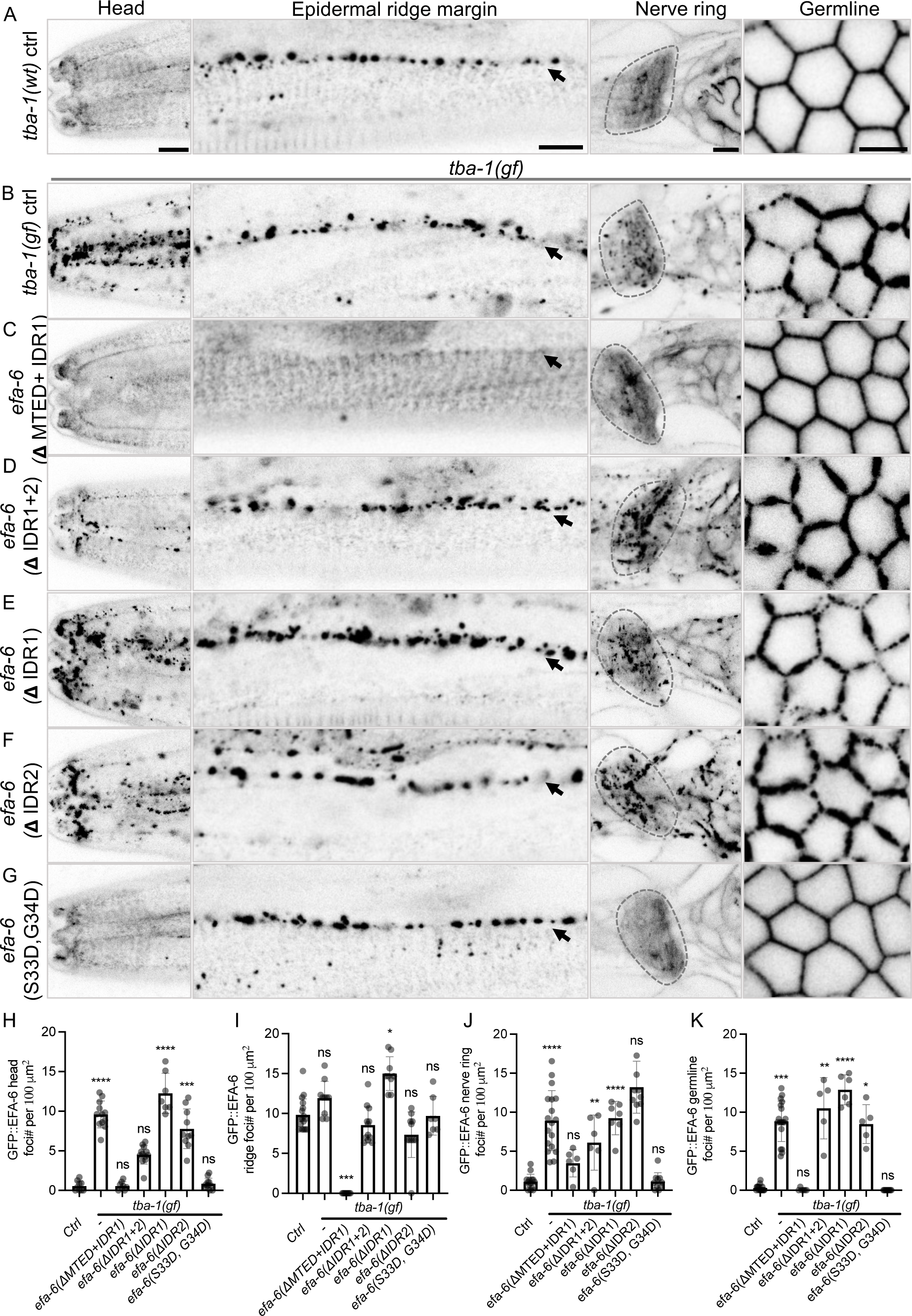
EFA-6 MTED is necessary for GFP::EFA-6 condensate formation in *tba-1(gf)* (A-G) Confocal images of GFP::EFA-6 in the head, epidermal ridge margin, nerve ring, and germline in 1-day adult animals of genotypes indicated on the left. Images are single slice of 0.5 µm thickness, scale = 5 µm. (H-K) Quantitation of GFP::EFA-6 condensates (foci) in head (H), lateral epidermal ridge margin (I), nerve ring (J), and germline (K). N = 10-15; Error bar shows SEM; P value determined by Kruskal-Wallis test, ns (not significant) P > 0.05, * P ≤ 0.05, ** P ≤ 0.01, *** P ≤ 0.001, **** P ≤ 0.0001.

### TBA-1(R241C)-induced EFA-6 condensates are stable compared to wild type and recruit TAC-1

To assess the dynamic properties of EFA-6 condensates in control and *tba-1(gf)* animals, we conducted fluorescence recovery after photobleaching (FRAP) analysis (Figure 7A, B). In control animals, GFP::EFA-6 condensates at the epidermal ridge margin recovered 70.5 ± 7.9 % (mean ± SEM, n = 9) of signal over 15 min. In contrast, GFP::EFA-6 condensates at the same region in *tba-1(gf)* animals recovered 20.5 ± 2.7 % of signal in 15 min (n = 17). We further performed FRAP analysis in *tba-1(0)* animals and observed recovery similar to the control. This analysis suggests that *tba-1(gf)* induced EFA-6 condensates display significantly lower EFA-6 turnover than in the wild type. Correlating with their reduced turnover, we observed that *tba-1(gf)* induced EFA-6 condensates tended to be more variable in shape than in the wild type, varying from spherical to elongated (Figure S6A).

**Figure 7.**
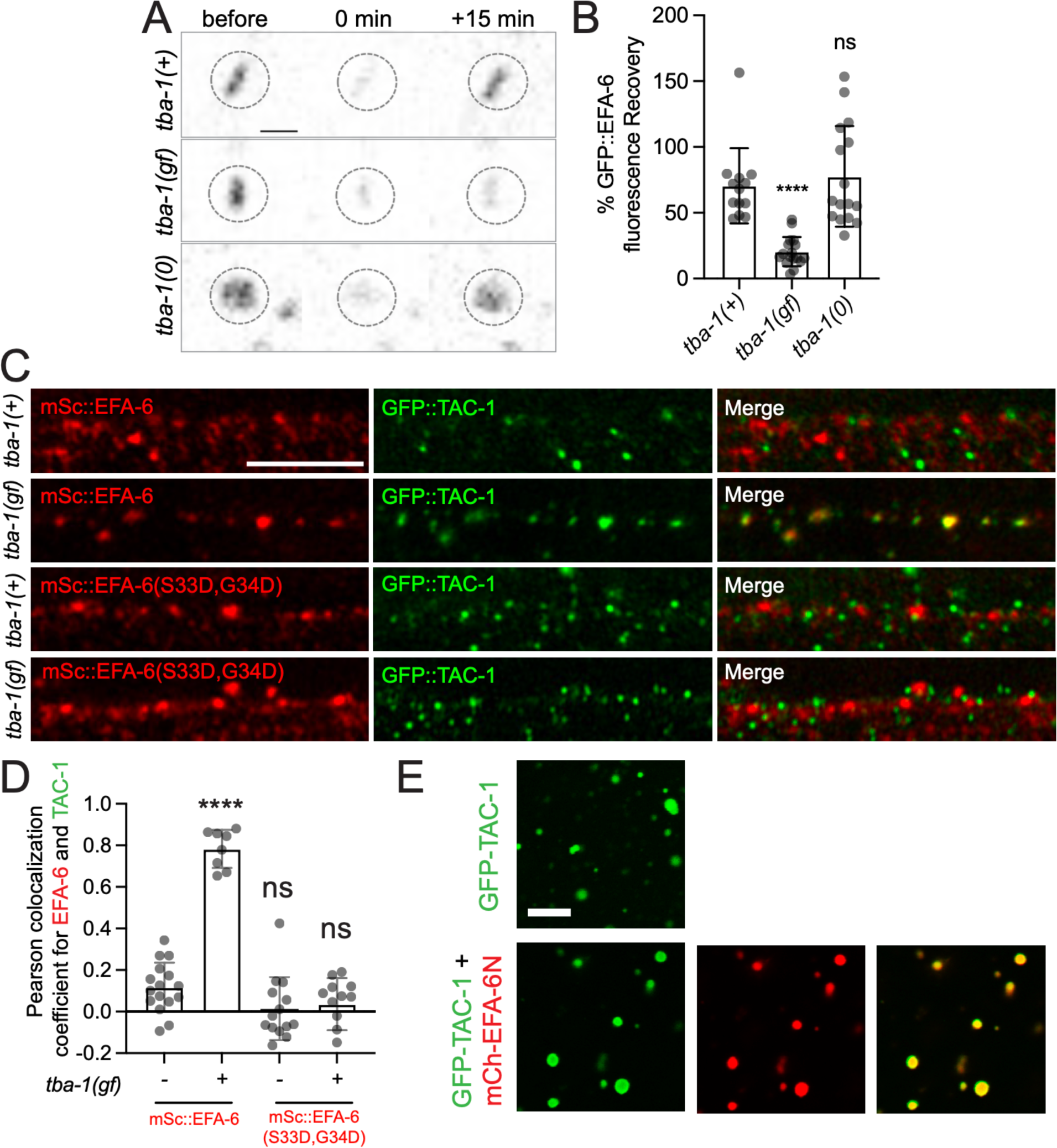
*tba-1(gf)* induced mSc::EFA-6 condensates display reduced turnover and recruit TAC-1 via the EFA-6 MTED. (A) Fluorescence recovery after photobleaching (FRAP) of epidermal GFP::EFA-6 condensates with genotypes indicated. Representative images show GFP::EFA-6 condensates before bleaching (t = −1 min), immediately after photobleaching (t = 0 min), and after recovery (t = 15 min); Scale = 1 µm. (B) Quantitation of percentage of GFP::EFA-6 condensates fluorescence recovery after 15 min (N = 6-7, two datapoint per animal). Statistics, Kruskal-Wallis test. Bar shows mean and SEM; ns, not significant; ****, P ≤ 0.0001. (C) Airyscan images show colocalization of GFP::TAC-1 with mSc::EFA-6(+) and mSc::EFA-6(S33D, G34D) in control *tba-1(+)* and *tba-1(gf)* mutant animals in the epidermal ridge margin. EFA-6 colocalizes with GFP::TAC-1 in *tba-1(gf)* but not in *tba-1(+)*; and EFA-6(S33D, G34D) does not show colocalization with GFP::TAC-1. All images are single slices 0.15 µm thickness; N = 6 per genotype. Scale = 5 µm. (D) Pearson colocalization coefficients of mSc::EFA-6 and GFP::TAC-1 in head epidermis. Statistics, Ordinary one-way ANOVA and Dunnett’s post test; **** P ≤ 0.0001. (E) EFA-6N and TAC-1 co-condensates *in vitro*. Representative images of droplets formed by GFP::TAC-1 (20 µM) alone and by mCherry::EFA-6N and GFP::TAC-1 (20 µM each) in droplet formation buffer. Scale = 10 µm.

TAC-1/TACC is a conserved component of pericentriolar material (Bellanger et al., 2007) that can bind to EFA-6 and colocalizes to axonal puncta in neurons after axon injury (Chen et al., 2015). We therefore examined the colocalization of mSc::EFA-6 and GFP::TAC-1 in wild type and in *tba-1(gf)*. In wild type control animals GFP::TAC-1 puncta formed in many tissues including the epidermis and these did not colocalize with mSc::EFA-6 condensates at the epidermal ridge margin (Figure 7C, D & S7). In contrast, in *tba-1(gf)* mutants, GFP::TAC-1 significantly colocalized with ectopic mSc::EFA-6 condensates, including in anterior epidermal cells (hyp5), nerve ring, and the syncytial germline (Figure 7C, D & S7). These EFA-6-colocalized TAC-1 puncta were consistently larger than TAC-1 puncta observed in the wild type. We further examined the colocalization of MT shaft marker MAPH-1.1 and minus-end binding proteins TBG-1 and GIP-2 with EFA-6 condensates in wild type control and *tba-1(gf)* mutants and found that they did not colocalize in either case (Figure S6). These data indicate that TAC-1 is specifically recruited to the ectopic EFA-6 condensates in *tba-1(gf)* mutants.

To address whether TAC-1 could co-condense with EFA-6 *in vitro,* we expressed and purified TAC-1 and performed *in vitro* droplet formation assays alone and with EFA-6N. We found that GFP::TAC-1 formed condensates alone and consistently co-condensed with mCh::EFA-6N (Figure 7E), supporting that TAC-1 and EFA-6 might interact in dynamic biomolecular condensates.

To evaluate if EFA-6 condensates might recruit TAC-1 in *tba-1(gf)* mutants, we examined GFP::TAC-1 localization in *efa-6(0)* mutants (Figure S8A). We found that *efa-6(0)* did not affect the overall pattern of TAC-1 puncta in *tba-1(+)* control but eliminated the larger TAC-1 puncta at epidermal ridge margins in *tba-1(gf)* mutants (Figure S8A, B). Conversely, to test whether TAC-1 could be involved in EFA-6 condensation, we examined a temperature-sensitive loss of function allele of *tac-1.* At the permissive temperature (20 °C), *tac-1(lf)* did not affect EFA-6 condensate formation in *tba-1(+)* animals or *tba-1(gf)*. However, at the restrictive temperature (25°C), *tac-1(lf)* partially reduced ectopic EFA-6 condensate formation in *tba-1(gf)* mutants (Figure S8C-E). These results suggest that TAC-1 is not absolutely required for ectopic EFA-6 condensate formation in the epidermis of *tba-1(gf)* animals but may play a role in promoting or stabilizing condensate formation.

Since EFA-6 MTED is required for *tba-1(gf)* induced GFP::EFA-6 condensates, we generated the same MTED mutation in mSc::EFA-6 animals. We found that in *tba-1(gf)* mutants, mSc::EFA-6(S33D, G34D) did not form ectopic condensates nor displayed colocalization with GFP::TAC-1 puncta at the epidermal ridge margin (Figure 7C, D). This suggests that the EFA-6 MTED is required for forming ectopic condensates in *tba-1(gf)* and is further required for recruitment of TAC-1 to the ectopic EFA-6 condensates.

## DISCUSSION

EFA6 proteins localize to the cellular cortex, where they have been shown to regulate MT dynamics in multiple cell types. By dissecting the localization of endogenous EFA-6, we uncovered a role for its intrinsically disordered region in forming previously undescribed cortical foci. Using an *in vitro* droplet assay we find that the EFA-6 N-terminal region is sufficient to form phase-separated condensates. Using forward genetics, we further reveal a critical role for α-tubulin in the regulation of EFA-6 localization across cell types. Our findings reveal intricate interactions between tubulins and a localized MT regulator in the cell cortex.

Previous studies of *C. elegans* EFA-6 relied on cell-type specific transgenic expression (Chen et al., 2015; O’Rourke et al., 2010). Our analysis of endogenously tagged EFA-6 shows that EFA-6 is expressed in nearly all types of cells and generally is localized to the cell cortex, consistent with the role of the PH domain in membrane association of EFA6 family members (Derrien et al., 2002). In addition, epidermal EFA-6 forms distinct cortical foci that are dependent on its N-terminal intrinsically disordered region (IDR). We find EFA-6 lacking the PH domain also forms punctate foci in response to heat stress. Moreover a 150 aa region of the EFA-6 N-terminus, including most of IDR1, is sufficient to form biomolecular condensates *in vitro*. Taken together these findings suggest that while EFA-6 is predominantly cortically localized, it can form condensates under specific conditions or stresses. As IDRs can promote phase separation (Uversky, 2017), we speculate the cortical foci of EFA-6 in the wild type epidermis and under heat stress may result from protein phase separation.

In *C. elegans,* most epidermal nuclei, organelles, and cytoplasm reside in thickened ridges whose margins are defined by body muscle quadrants (White et al., 1986). Wild type EFA-6 foci gradually form at the cortex of sublateral ridge margins, raising the question of the cues that localize EFA-6 foci. Sublateral ridge margins also contain stable colchicine-insensitive MT bundles (Chuang et al., 2016; Wang et al., 2015), as well as MT plus and minus end binding proteins distributed in punctate patterns (Castiglioni et al., 2020). Speculatively, EFA-6 cortical foci in these regions may be related to the presence of specific types of MTs. The MTED of other EFA6 proteins can inhibit MT polymerization *in vitro* (Qu et al., 2019). We find EFA-6 cortical foci do not colocalize with but are in proximity to MT plus and minus end markers. These findings might be reconciled if a transient interaction between EFA-6 and tubulin can inhibit EFA-6 condensation and locally destabilize MTs. Overall, these findings indicate epidermal sublateral ridge margins contain complex subcellular compartments.

The evidence for a specific interaction between tubulins and EFA-6 is bolstered by our finding, through unbiased genetic screening, that a gain of function in α-tubulin alters EFA-6 cortical foci formation. We also show an equivalent change in *tba-2* affects EFA-6 condensation in a similar manner, indicating these effects are specific to α-tubulin. TBA-1 and TBA-2 are widely coexpressed and functionally redundant α-tubulins (Nishida et al., 2021; Phillips et al., 2004). TBA-1(gf) caused the formation of ectopic EFA-6 foci in all cell types examined, ranging from neurons to epidermis to undifferentiated embryonic blastomeres. Thus, in most cell types, the cortical localization of EFA-6 depends on normal α-tubulin function.

The EFA-6 foci formed in *tba-1(gf)* appear to be distinct from those seen in the wild type in several ways. First, *tba-1(gf)*-induced EFA-6 foci are seen in almost all tissues examined, whereas in the wild type EFA-6 punctate foci are restricted to margins of epidermal ridges. Second, based on FRAP experiments TBA-1(gf) induced EFA-6 foci show reduced turnover compared to wild type, suggesting they are more stable. Third, *tba-1(gf)*-induced EFA-6 foci require the MTED, whereas wild-type EFA-6 foci depend on its IDR. Fourth, *tba-1(gf)* induced EFA-6 foci recruit TAC-1. TAC-1 was previously shown to bind the EFA-6 N-terminus and to colocalize with EFA-6 at injury-induced axonal puncta in neurons (Chen et al., 2015). Here, we find that purified TAC-1 and the EFA-6 N-terminus can co-condense *in vitro*. TAC-1 forms puncta in the epidermis that do not colocalize with EFA-6 cortical foci in the wild type. In *tba-1(gf)*, EFA-6 recruited TAC-1 to ectopic cortical foci, dependent on the EFA-6 MTED. TAC-1 was not absolutely required for EFA-6 condensate formation in wild type or in *tba-1(gf)*, however EFA-6 foci appeared smaller and less dense in *tac-1(lf)* mutants. This evidence suggests a role for EFA-6 in relaying signals from tubulins to the cortical cytoskeleton and TAC-1.

The organization of MT arrays in most tissues was largely normal in *tba-1(gf).* However, TBA-1(gf) itself displayed reduced incorporation into MTs. In *tba-1(gf),* ectopic EFA-6 foci might be triggered by unincorporated TBA-1(gf) or aberrant TBA-1(gf)-containing MTs. Formation of the aberrant TBA-1(gf) foci was suppressed by loss of function in *tbb-2/*β-tubulin. Moreover, loss of function in the tubulin chaperone *evl-20*/Arl2 also reduced EFA-6 foci formation by TBA-1(gf). Loss of function in *evl-20* reduces organized MTs in embryos (Antoshechkin and Han, 2002). In view of these findings and the dominant nature of *tba-1(gf)*, we speculate that an abnormal TBA-1(gf)/TBB-2 heterodimer causes ectopic EFA-6 condensation.

The R241 residue is invariant in all α-tubulins and is located at the C-terminal end of helix H7. Unusually, the charged R241 side chain is fully buried, making H-bonds to several other residues. Our analysis indicates the R241 side chain is critical for function, as missense changes to C, A, K, or E all had deleterious effects; mutation of R241 to K or E caused fully penetrant lethality, as did R241C in TBA-2. Our recovery of R241C as a viable mutant suggests this mutation causes a moderate gain of function. There has been little mechanistic analysis of R241 in other systems, although in *Toxoplasma* α-tubulin R241C confers resistance to dinitroaniline inhibitors (Morrissette et al., 2004) and some human cancers contain SNPs affecting α-tubulin R241 (Abbaali et al., 2023). It is not straightforward to predict the effects of α-tubulin R241 mutations from available MT structures, and it will be important to examine the effects of R241 mutation in future structural or biophysical studies. Our finding that *efa-6(0)* can suppress *tba-1(gf)* lethality suggests that the deleterious effects of tubulin gain of function *in vivo* may in part be mediated by EFA-6.

Mammalian EFA6 family members have been linked to epithelial polarity (Luton et al., 2004), tight junction formation (Klein et al., 2008), endocytosis (Boulakirba et al., 2014), apical luminogenesis (Milanini et al., 2018), and ciliogenesis (Partisani et al., 2021). So far, most studies of EFA6 members in epithelia have focused on their Arf6 GEF activity. In *C. elegans* loss of *efa-6* function does not dramatically affect epidermal development, suggesting EFA-6 might act redundantly with other factors in epithelial development. A parallel may be drawn with the MT regulator PTRN-1/Patronin, which is not essential for epidermal development but whose function is revealed in sensitized backgrounds (Chuang et al., 2016; Wang et al., 2015). In neurons, EFA-6 function becomes critical during responses to axon injury (Chen et al., 2015), and it will be of interest to examine whether EFA-6 functions in epithelial cytoskeletal responses to wounding (Taffoni et al., 2020). Overall our findings reveal a complex interplay between the MT cytoskeleton and cortical regulators that may become critical in certain stress conditions or subcellular locations.

## METHODS

### C. elegans genetics

All strains of *C. elegans* were maintained on nematode growth media (NGM) plates seeded with *E. coli* OP50 at 20 °C, except where noted. Compound mutants with desired fluorescent reporters (KI or transgenes) were generated by genetic crosses; genotypes were confirmed using allele-specific polymorphisms or visualization of fluorescence. Key reagents, including strains and oligonucleotides, are described in Supplemental Table 1. Details of allele detection are in Supplemental Table 2. Sequences of CRISPR reagents and edits are in Supplemental Table 3.

### CRISPR-Cas9 genome editing

#### GFP::EFA-6(ju1658) knock-in

We generated GFP::EFA-6(*ju1658*) knock-in using the CRISPR-Cas9 Self-Excising Cassette (SEC) method (Dickinson et al., 2015). The sgRNA 5’-CGTTTTCAGAGTGATGGCGA-3’ targeting the *efa-6* ATG was selected using CRISPR design tool (http://crispr.mit.edu) and was inserted into pDD162 vector (Addgene #47549) using primers YJ12362 and YJ12363 (Table S2/3) to generate pCZ986. To generate the homologous repair template clone pCZ996, ∼1 kb DNA flanking either side of the target sequence was amplified using primers YJ12364 and YJ12365, YJ12369 and YJ12370, respectively, and cloned into pDD282 (Addgene #66823, FP::SEC vector) by Gibson assembly (NEB). A DNA mix containing pCZ986 (50 ng/µL), pCZ996 (10 ng/µL), co-injection marker pCFJ104 (P*myo-3*-mCherry; Addgene #19328, 5 ng/µL) and marker pCFJ90 (P*myo-2*-mCherry; Addgene #19327, 2.5 ng/µL) was injected into N2 young adult hermaphrodites. Progeny were treated with hygromycin and surviving animals that showed the Rol phenotype but lacked co-injection marker expression were selected as candidate knock-in GFP-SEC-EFA-6(*ju1655*). GFP-SEC-*efa-6*(*ju1655*) animals were heat shocked at 34 °C for 4 h to excise the SEC, resulting in GFP::EFA-6(*ju1658*). Sequencing analysis of both genomic DNA and RT-PCR products from *ju1658* confirmed that GFP was inserted in-frame at the ATG of the EFA-6A, -B, and -C isoforms, but not the D isoform. *mSc::EFA-6(syb6998)* was obtained from SunyBiotech (Fuzhou, China), with mScarlet coding sequence inserted at the exact same nucleotide as in *ju1658.* Primers used for plasmid generation are listed in Table S1 and sequencing primers in Table S2.

#### Genome editing of GFP::EFA-6(ju1658), tba-1, tba-2

Genome editing was done following the melting method as described (Ghanta and Mello, 2020). sgRNAs were selected using IDT database and are listed in Table S3. 1-5 μM of target crRNAs were incubated with 20 μM tracrRNA (IDT) at 95°C for 5 min in a volume of 5 μL. The reaction was cooled to RT for 5-10 min, and 2 μM of Cas9 protein (Macrolabs, UC Berkeley) was added and incubated at 25°C for 1-2 h. Homology repair templates were designed to create the desired mutation and silent mutations in the crRNA recognition sequence, PAM, as well as an enzyme digestion site to assist genotyping of edits. Homology repair templates of ∼200 bp (5 μM, IDT) used for SNP editing, were added to the crRNA-tracrRNA-Cas9 mix and injected into the animals with 0.5-1 μM of pCFJ90 (P*myo-2*-Cherry) as a co-injection marker. About 15 P_0_ animals were injected in each experiment and transferred to 5 seeded NGM plates (2-3 animals/plate) to propagate for 3-4 days at 20°C. From plates where co-injection marker was observed, multiple F_1_ progeny were isolated and screened for correct editing using PCR coupled with enzyme digestion. All edited alleles were outcrossed to N2 at least four times; sequence changes and genotyping information are in Tables S2 and S3.

### Mutagenesis screen for altered localization pattern of GFP::EFA-6(*ju1658*)

To identify genes that regulate GFP::EFA-6 expression or localization we mutagenized CZ26544 (GFP::EFA-6(*ju1658*); P*mec-7*-mRFP (*jsIs973*)) with 47 mM Ethyl Methane Sulphonate (EMS) for 4 h following standard procedures (Brenner, 1974). P_0_ animals were washed with M9 buffer, and after 1 h recovery at room temperature, 32 healthy-looking P_0_ animals were placed on 8 plates to produce F_1_ progeny. About 1,080 F_1_ animals were transferred to freshly seeded plates in groups of 3 to produce F_2_ progeny. Among F_2_ progeny from each F_1_ plate, 20-30 L4 animals were mounted on 5% agarose pad with 2 mM levamisole and observed under Zeiss Axioplan 2 microscope equipped with Chroma HQ filters and a 63x objective. Animals showing significant changes of GFP::EFA-6 pattern were recovered to propagate and the phenotype verified in the F_3_ generation. One F_2_ line per F_1_ plate was kept to ensure independent isolates. Mutants were outcrossed to CZ27027 GFP::EFA-6(*ju1658*) and reisolated based on altered GFP::EFA-6 phenotypes using compound microscope.

### Whole genome sequencing analysis of *tba-1(ju1761)*

We obtained WGS data from the parent strain CZ26544 (GFP::EFA-6(*ju1658))* and two outcrossed versions of *ju1761,* strains CZ27301 (GFP::EFA-6(*ju1658; ju1761*) and CZ27771 (GFP::EFA-6(*ju1658; ju1761*). Genomic DNAs were prepared using a Gentra Puregene Kit (Qiagen). 20x coverage whole-genome sequence (WGS) was obtained as 90 bp paired-end reads (BGI Americas). We mapped the raw reads to the *C. elegans* reference genome (WBcel235/ce11) using the Galaxy platform (http://usegalaxy.org). We then used MiModd 0.1.9 (https://mimodd.readthedocs.io/ed/latest/#) to subtract mutations present in the parental strain (CZ26544), leaving candidate induced mutations. We analyzed the EMS-induced mutations that co-segregated with *ju1761* phenotype and identified genetic linkage to Chromosome I. *ju1761* contains a C>T transition (CGT > TGT) in *tba-1*, resulting in Arg241Cys missense mutation. Additional outcrossing of *ju1761* was conducted to eliminate linked EMS-induced SNPs prior to analysis. We also edited the same nucleotide change in N2 background as described above, designated *ju1869*, which exhibited mutant phenotypes indistinguishable from *ju1761* and was used in most strains (Table S1).

### Fluorescence microscopy

Animals were paralyzed using 5 mM levamisole in M9 buffer on 4% agar pad. Standard confocal fluorescence imaging was performed on a Zeiss LSM800 (Axio Observer.Z1/7) using a Planapochromat 63x/NA1.4 oil immersion objective, with pinhole sizes at 100 μm, 1.44-3.08 Airy units, laser power of 1% for 488 nm and 10% for 561 nm. Detector gain was 678 V for GFP and 750 V for mScarlet. Z-stack images were acquired with 0.5 μm interval; images were analyzed using Image J and single slices shown as specified in legends. For imaging MT markers (tubulins or MAPH-1.1) in epidermis, young adult animals were paralyzed in 0.5 mM levamisole in M9 buffer on 4% agar pad, which took ∼30 min; images were acquired within the next 30 min on Zeiss confocal LSM800 microscope using 100x/NA1.4 oil immersion objective, 1% 488 nm laser power, and default detector gain and gain of 680 V and 1, respectively.

For colocalization analysis of GFP::EFA-6 or mSc::EFA-6 with MT or other epidermal protein markers, we used Zeiss LSM900 equipped with Airyscan (Axio Observer.Z1/7) in SuperResolution mode with ‘Best Signal’ detection setup to prevent laser bleed through. GFP::LAM-1 (*qyIs8*) was imaged using a Planapochromat 63x/NA1.4 oil immersion objective and 2.5% 488 nm laser power. GFP tagged MAPH-1.1, EBP-2, TAC-1, GIP-2, and mCherry tagged TBG-1 were imaged using a 100x/NA1.46 immersion oil objective, with 4.5% 488 nm laser power (EBP-2, TAC-1, and GIP-2) and 5% 561 nm laser power (TBG-1), respectively. mSc::EFA-6 was imaged with 5% 570 nm laser power, and GFP::EFA-6 was 2.5% 488 nm laser power. Z-stacks were acquired with 0.15 μm interval for EBP-2, TAC-1, and GIP-2, and 0.13 μm interval for TBG-1 and MAPH-1.1. Detector gain and detector digital gain was set to default of 850 V and 1, respectively.

### Quantitative image analysis of GFP::EFA-6 and microtubules

For quantitative analysis of GFP::EFA-6 cortical foci, we collected confocal z-stacks spanning the lateral epidermal ridge with 0.5 μm intervals and analyzed individual z sections using ImageJ. We set the threshold for each image to capture all fluorescent foci as scored manually, and calculated number using Analyze Particles function with size threshold of 0.1 μm^2^. Number of puncta per 100 μm^2^ was deduced from each ROI [(No. of puncta per ROI/Area of ROI) *100] and plotted using Prism 9. ROIs were selected differently for each tissue: in the head epidermis, pentagonal ROIs roughly outlined hyp5 cell; for lateral epidermis ridge margin, ROIs were 5 x 60 μm rectangles containing the ventral ridge margin; nerve ring ROIs are outlines of the lateral neuropil; for germline, 100 μm^2^ ROIs were randomly selected outlining the edges of the ‘honeycomb’ pattern in the syncytial germline. For germline, a combination of manual and quantitative method was used to count foci. If no foci were observed, images were recorded as zero without thresholding. Images containing one or more foci were subjected to a threshold for quantitation. For quantitating large GFP::TAC-1 puncta in Figure S8, we set an area threshold of 0.13 μm^2^ (mean wild type GFP::TAC-1 puncta area) and measured the number of puncta above the threshold in three independent ROI of 15 x 3 μm along the lateral ridge margin.

To measure mean intensity of GFP::EFA-6 foci along epidermal ridge margin, rectangular ROIs of 2 x 13 μm were selected and measured in ImageJ. 2-3 ROIs were selected in each animal, mean intensity per 100 μm^2^ was deduced and plotted as described above. Mean intensity of GFP::EFA-6 in different tissues in Figure S2 were quantified similarly in Image J: for nerve ring, ROIs were selected outlining the whole neuropil, one per animal; for germline, 6 linear ROIs were selected randomly along the cell membrane for each animal and the mean shown as one data point per animal; for epidermis, rectangular ROIs were selected at the lateral anterior epidermis region, three data points per animal.

For analysis of circumferential microtubule bundles in the epidermis, visualized with tubulin or MAPH-1 markers, we obtained single plane confocal images, which were saved as .czi files and analyzed using the ‘Measure’ option in ImageJ Analysis menu. For MT bundle intensity, we drew ROI as a line along the circumferential bands and measured the mean intensity along the ROI using ImageJ. Mean intensity for each line ROI was normalized to the length of the ROI to calculate mean intensity 1 μm per length. To count MT bundles, we drew 20 μm lines orthogonal to circumferential bands near both lateral epidermal ridges in an image, then using profile plot counted number of MT bundles along the length of line ROI. Each profile plot peak was considered as an independent MT bundle.

For quantitative analysis of colocalization (e.g. of EFA-6 and TAC-1), we analyzed a single plane image from Airyscan .czi stacks of 0.15 μm interval. We drew two ROIs, each of 100 μm^2^, in the hyp4 and hyp5 region of the head. Using the colocalization tool in Zeiss Zen software, we obtained the Pearson’s colocalization coefficient for each ROI. To analyze TAC-1 and EFA-6 colocalization profiles at the ridge margin, a single plane image from the .czi stack was converted to an RGB image using ImageJ. A line ROI of 10 μm along the length of ridge was drawn and line scan profiles were obtained using the RGB profiler plugin in ImageJ.

### Fluorescence recovery after photobleaching (FRAP)

FRAP of GFP::EFA-6 was performed using a Zeiss LSM900 microscope in confocal mode. Circular regions of interest (ROIs) for acquisition and photobleaching of GFP::EFA-6 were set in the epidermis and then bleached with 7 iterations of 100% 488 nm laser power. After bleaching, recovery was monitored at 5 min intervals for 15 min. A nearby condensate was used as control and unbleached non-condensed signal used to subtract background. Mean fluorescence intensity after bleaching was calculated as percentage of the unbleached ROI intensity.

### RNA interference

For systemic RNA interference by feeding (Kamath et al., 2003), we grew HT115(DE3) containing empty vector control or *tba-1* dsRNA (RNAi clone F26E4.8) in LB broth with 50 µg/ml carbenicillin overnight at 37 °C. Bacteria were plated on NGM plates containing 5 mM isopropyl-D-thiogalactoside (IPTG) and 50 µg/ml carbenicillin. Seeded plates were kept at 25 °C for 24 h. Ten gravid adults were allowed to lay eggs for 2-3 h, and then removed. Progeny were maintained at 20 °C and day 1 adults were imaged on compound microscope.

### GFP::EFA-6 *trans*-heterozygote imaging

10-15 GFP::EFA-6; P*mec-7*::RFP males were crossed with five GFP::EFA-6; *tba-1(gf)* hermaphrodites at 20 °C. 10-15 L4 hermaphrodite cross progeny were transferred to a fresh plate. 10-15 GFP::EFA-6 with homozygous *tba-1(+)* or *tba-1(gf)* L4 animals were also isolated on the same day as controls. Animals were maintained at 20 °C and imaged after 24 h using the compound microscope. For imaging GFP::TBA-1(+) *trans*-heterozygotes with *tba-1(+)* and *tba-1(gf),* 10-15 GFP::TBA-1(+) homozygous males were crossed with 5-6 N2 or *tba-1(gf)* hermaphrodites. 10-15 L4 hermaphrodite F_1_ progeny expressing GFP::TBA-1 were transferred to fresh OP50 plates, and maintained at 20 °C. After 24 h, gravid adults were imaged using LSM800 confocal microscope.

### Temperature shift of *tac-1(or455)*

*tac-1(or455)* mutants display temperature sensitive embryonic lethality and resemble null mutants when raised at the restrictive temperature (Bellanger et al., 2007). We maintained *tac-1(or455)-*containing strains at 15°C. We synchronized animals by picking 10 adults to a freshly seeded plate and allowing to lay eggs for 2.5 h at 15 °C; adults were removed and progeny maintained at 15°C for 60 h. Plates were transferred to 25 °C for 36 h before imaging on LSM800.

### Neuronal morphology scoring

We scored PLM morphology in day 1 adults using the touch neuron marker P*mec-7*-mRFP(*jsIs973*) (Zheng et al., 2011). Scoring was conducted over multiple days in genotype-blind manner. PLM was scored as overshooting if the tip of the process was anterior to the midline of ALM soma (Chen et al., 2011).

### Heat stress experiment

L4 animals were maintained at 20 °C and day 1 adults incubated at 34 °C for 2 h or kept at 20 °C as control. Immediately after heat stress, animals were immobilized with 10 mM levamisole in M9 buffer on 4% agar pad. Images were taken with Zeiss LSM900 equipped with Airyscan (Axio Observer.Z1/7) in SuperResolution mode. To score condensate recovery after heat stress, mixed stage animals were treated with heat stress at 34 °C for 2 h. ∼30 treated or control adults were immobilized with 5 mM levamisole on 4% agar pad and visualized using Zeiss Axioplan 2 microscope equipped with Chroma HQ filters and a 63x objective. Animals were observed every 2 h for 8 h, and number of animals showing condensates was counted.

### Embryonic lethality assay

10-15 healthy L4 animals cultured at 20 ℃ were transferred to fresh plates, shifted to 25 °C as L4s then maintained at 25 ℃. Each animal was transferred to a fresh plate every 24 h for 3 days. Hatched progeny were counted on the 3rd day after transfer. Animals were immobilized with 0.1 mg/ml NaN_3_ on NGM agar plates to facilitate counting.

### Protein analysis

To synchronize animal culture, 15-20 gravid adults were placed on a NGM plate seeded with OP50 to lay eggs for 8-10 h and removed. Progeny were maintained at 20°C for three days then collected using M9 buffer in a 15 ml falcon tube (no centrifuge) under gravity. After washing with M9 buffer twice, excess M9 buffer was removed to a final volume of 1 ml, and packed worms were transferred to 2 ml Eppendorf tube using a glass pipette. Excess M9 buffer was removed to leave about 300-500 µl of loose worm pellets. An additional quick wash was performed with 1 ml of lysis buffer (50 mM Tris HCl, 150 mM NaCl, 10% glycerol, 0.1% Tergitol, PhosphoSTOP, CompleteO, 5 mM NaF, 1 mM Na_2_MoO_4,_ and 3 mM PMSF) (Crawley et al., 2019), and worms were pelleted at 2348 rcf for 2-3 min. Fresh lysis buffer was added to worm pellets in 3:1 (volume), and samples stored at −80 °C. To make protein extracts, frozen worm pellets were thawed on ice and refrozen in pre-chilled mortar and pestle in liquid N_2_. After grinding, worm powders were collected in 2 ml Eppendorf tubes and centrifuged at 21130 rcf at 4 °C for 30 min. Supernatants (about 200-250 µl) were collected into fresh Eppendorf tubes, and protein concentration was quantified using Pierce^TM^ BCA protein assay kit and NOVOstar (BMG labtech). For western blotting, 20-30 µg of proteins in 1x NuPage (Invitrogen) LDS sample buffer and NuPAGE sample reducing agent (15 µl final volume) was run on Invitrogen BOLT 4-12% gradient pre-casted gels using 1x Bolt^TM^ MES SDS running buffer in Invitrogen mini tank setup. Samples were transferred to nitrocellulose membrane (0.45 μm, BioRad) using 1x Pierce^TM^ Western Blot Transfer buffer with 20% methanol (v/v, Fisher Chemicals, HPLC grade) in BioRad electrotransfer setup at a constant current of 100 mA for 90 min at 4°C. Membrane blots were incubated with primary antibody overnight at 4 °C, anti-FLAG (1:5000), anti-tubulin (1:5000), or anti-actin (1:10,000), and secondary antibodies (1:10,000, anti-rabbit for flag and anti-mouse tubulin and actin) at RT for 60-90 min then visualized using Image Studio Ver 5.2 and LI-COR ODYSSEY imaging system.

### Recombinant Protein Expression

The Rosetta BL21(DE3) *E. coli* strain was used for recombinant protein expression with pET expression vectors. Single colonies from an overnight culture were used to inoculate a 5 mL culture volume of LB and antibiotics and 1% glucose which was incubated overnight at 37 °C shaking at 225 rpm. Small volume overnight cultures were then used to inoculate >200 mL culture volume of LB, antibiotics, and 1% glucose. The inoculated culture was incubated at 37 °C with shaking at 225 rpm until OD_600_ was between 0.4 and 0.7. When the appropriate culture density was achieved, the culture media was then cooled on ice prior to induction of protein expression by addition of 0.1 mM IPTG. The induced culture was further incubated at 18 °C with shaking at 120 rpm overnight. Following this, the culture was pelleted by repeated centrifugation at 4 °C, 5000 rpm for 10 min. Bacterial pellets were stored at −80 °C for at least 30 min prior to protein isolation by affinity chromatography.

### Affinity Chromatography

Induced bacterial pellets were placed on ice, resuspended in 15 mL of Buffer A (50 mM Tris pH 7.5, 500 mM NaCl, 10 mM Imidazole) and lysed by sonication on ice. Sonication proceeded by pulsing at 30% amplitude for 1 s on 5 s off for 12 min total run time. Lysed bacterial suspensions were sedimented by centrifugation at 18,000 *g* for 30 min at 4 °C in a Sorvall 2B ultracentrifuge with appropriate rotor. The supernatant was transferred to a clean 50 mL Falcon tube with 1 mL of pre-equilibrated, with Buffer A, NiNTA agarose beads for binding for 1.5 h at 4 °C with light agitation. The NiNTA agarose beads with bound protein were then loaded onto BioRad manual chromatography columns and allowed to sediment by gravity prior to washing with 15 mL of Buffer A. The bound protein was then eluted with Elution Buffers (EB) 1-3 containing a stepped gradation of 50 mM, 100 mM, and 250 mM Imidazole. The protein of interest and purity was confirmed within elutions by SDS-PAGE and Coomassie Blue staining prior to combining protein containing elutions into a 10k MWCO dialysis tube and dialysis overnight in 1L Dialysis Buffer (50 mM Tris pH 7.5, 125 mM NaCl, 10% Glycerol, 1 mM DTT).

### Droplet Formation Assays

After dialysis, concentration of purified recombinant proteins was measured, followed by dilution using dialysis buffer to desired concentration. The protein solution was mixed with the droplet formation buffer (Dialysis buffer with 20% or 40% m/v PEG-8000) at 1:1 ratio prior to imaging on a Zeiss LSM780 scanning confocal microscope. All comparisons utilized the same concentration of molecular crowding agents with protein concentration as stated. Droplet size quantitation was performed in Fiji.

### Statistical Analysis

All statistical analysis used GraphPad Prism 9. Data passing normality tests were analyzed by parametric tests such as ordinary one-way ANOVA and an appropriate post test. Data not passing the normality test were analyzed using nonparametric tests i.e. Mann-Whitney test for comparing two data sets or Kruskal-Wallis test for > 2 comparisons.

## Supplemental Materials: 8 Supplemental Figures and 3 Supplemental Tables

Table S1. Key resources

Table S2. Allele information and detection methods

Table S3. CRISPR sgRNA and edited sequences

## ACKNOWLEDGEMENTS

We thank members of our labs for discussion and advice during this work. We thank Steve Blazie for advice on biochemistry, and Kancheng Yin and Jianing Shi for constructing strains. We appreciate insightful discussion with Andres Leschziner and Luke Rice on tubulins, and Andreas Ernst for comments on the manuscript. Some strains were supplied by the *Caenorhabditis* Genetics Center (CGC), which is funded by NIH P40 OD10440. Supported by NIH grants NIA R01AG070214 and R01AG071591 to LC, R01NS093988 to YJ and ADC, R35NS127314 to YJ, and R35GM134970 to ADC.

## Author Contributions

GFP::EFA-6 was generated by XM and NHT. The genetic screen was performed by XW and NHT. AS and XL generated *efa-6* alleles, performed imaging, and conducted genetic interaction studies. AS performed mapping and cloning of *tba-1*, performed biochemistry and imaging. GG, INS and LC analyzed EFA-6 condensation *in vitro*. LC, YJ, and ADC secured funding and analyzed data. AS, XL, LC, YJ and ADC wrote the paper with input from other authors.

## Resource Availability

### Lead Contact

Further information and requests for resources and reagents should be directed to and will be fulfilled by the lead contact, Andrew Chisholm

### Materials availability

All unique materials generated in this study, including strains and plasmids will be shared by the lead contact upon request.

### Data and code availability

Key quantitative data will be supplied in a source data file. Key images will be deposited in the Figshare repository. Additional data reported in this paper are available from the lead contact upon request.

## REFERENCES

Abbaali, I., Truong, D., Day, S.D., Mushayeed, F., Ganesh, B., Haro-Ramirez, N., Isles, J., Nag, H., Pham, C., Shah, P., Tomar, I., Manel-Romero, C., Morrissette, N.S., 2023. The tubulin database: Linking mutations, modifications, ligands and local interactions. PLoS One 18, e0295279.

Al-Bassam, J., 2017. Revisiting the tubulin cofactors and Arl2 in the regulation of soluble alphabeta-tubulin pools and their effect on microtubule dynamics. Mol Biol Cell 28, 359–363.

Alushin, G.M., Lander, G.C., Kellogg, E.H., Zhang, R., Baker, D., Nogales, E., 2014. High-resolution microtubule structures reveal the structural transitions in alphabeta-tubulin upon GTP hydrolysis. Cell 157, 1117–1129.

Andrusiak, M.G., Sharifnia, P., Lyu, X., Wang, Z., Dickey, A.M., Wu, Z., Chisholm, A.D., Jin, Y., 2019. Inhibition of Axon Regeneration by Liquid-like TIAR-2 Granules. Neuron 104, 290–304 e298.

Antoshechkin, I., Han, M., 2002. The *C. elegans evl-20* gene is a homolog of the small GTPase ARL2 and regulates cytoskeleton dynamics during cytokinesis and morphogenesis. Dev Cell 2, 579–591.

Bellanger, J.M., Carter, J.C., Phillips, J.B., Canard, C., Bowerman, B., Gonczy, P., 2007. ZYG-9, TAC-1 and ZYG-8 together ensure correct microtubule function throughout the cell cycle of *C. elegans* embryos. J Cell Sci 120, 2963–2973.

Belmont, L.D., Mitchison, T.J., 1996. Identification of a protein that interacts with tubulin dimers and increases the catastrophe rate of microtubules. Cell 84, 623–631.

Boulakirba, S., Macia, E., Partisani, M., Lacas-Gervais, S., Brau, F., Luton, F., Franco, M., 2014. Arf6 exchange factor EFA6 and endophilin directly interact at the plasma membrane to control clathrin-mediated endocytosis. Proc Natl Acad Sci U S A 111, 9473–9478.

Brenner, S., 1974. The genetics of *Caenorhabditis elegans*. Genetics 77, 71–94.

Bu, S., Yong, W.L., Lim, B.J.W., Kondo, S., Yu, F., 2021. A systematic analysis of microtubule-destabilizing factors during dendrite pruning in Drosophila. EMBO Rep 22, e52679.

Castiglioni, V.G., Pires, H.R., Rosas Bertolini, R., Riga, A., Kerver, J., Boxem, M., 2020. Epidermal PAR-6 and PKC-3 are essential for larval development of *C. elegans* and organize non-centrosomal microtubules. Elife 9.

Chaaban, S., Jariwala, S., Hsu, C.T., Redemann, S., Kollman, J.M., Muller-Reichert, T., Sept, D., Bui, K.H., Brouhard, G.J., 2018. The Structure and Dynamics of *C. elegans* Tubulin Reveals the Mechanistic Basis of Microtubule Growth. Dev Cell 47, 191–204 e198.

Chakrabortee, S., Byers, J.S., Jones, S., Garcia, D.M., Bhullar, B., Chang, A., She, R., Lee, L., Fremin, B., Lindquist, S., Jarosz, D.F., 2016. Intrinsically Disordered Proteins Drive Emergence and Inheritance of Biological Traits. Cell 167, 369–381 e312.

Chen, L., Chuang, M., Koorman, T., Boxem, M., Jin, Y., Chisholm, A.D., 2015. Axon injury triggers EFA-6 mediated destabilization of axonal microtubules via TACC and doublecortin like kinase. Elife 4.

Chen, L., Wang, Z., Ghosh-Roy, A., Hubert, T., Yan, D., O’Rourke, S., Bowerman, B., Wu, Z., Jin, Y., Chisholm, A.D., 2011. Axon regeneration pathways identified by systematic genetic screening in *C. elegans*. Neuron 71, 1043–1057.

Chen, L., Zhang, Z., Han, Q., Maity, B.K., Rodrigues, L., Zboril, E., Adhikari, R., Ko, S.H., Li, X., Yoshida, S.R., Xue, P., Smith, E., Xu, K., Wang, Q., Huang, T.H., Chong, S., Liu, Z., 2023. Hormone-induced enhancer assembly requires an optimal level of hormone receptor multivalent interactions. Mol Cell 83, 3438–3456 e3412.

Chuang, M., Hsiao, T.I., Tong, A., Xu, S., Chisholm, A.D., 2016. DAPK interacts with Patronin and the microtubule cytoskeleton in epidermal development and wound repair. Elife 5.

Crawley, O., Opperman, K.J., Desbois, M., Adrados, I., Borgen, M.A., Giles, A.C., Duckett, D.R., Grill, B., 2019. Autophagy is inhibited by ubiquitin ligase activity in the nervous system. Nat Commun 10, 5017.

Derrien, V., Couillault, C., Franco, M., Martineau, S., Montcourrier, P., Houlgatte, R., Chavrier, P., 2002. A conserved C-terminal domain of EFA6-family ARF6-guanine nucleotide exchange factors induces lengthening of microvilli-like membrane protrusions. J Cell Sci 115, 2867–2879.

Dickinson, D.J., Pani, A.M., Heppert, J.K., Higgins, C.D., Goldstein, B., 2015. Streamlined Genome Engineering with a Self-Excising Drug Selection Cassette. Genetics 200, 1035–1049.

Eva, R., Koseki, H., Kanamarlapudi, V., Fawcett, J.W., 2017. EFA6 regulates selective polarised transport and axon regeneration from the axon initial segment. J Cell Sci 130, 3663–3675.

Fayad, R., Rojas, M.V., Partisani, M., Finetti, P., Dib, S., Abelanet, S., Virolle, V., Farina, A., Cabaud, O., Lopez, M., Birnbaum, D., Bertucci, F., Franco, M., Luton, F., 2021. EFA6B regulates a stop signal for collective invasion in breast cancer. Nat Commun 12, 2198.

Franco, M., Peters, P.J., Boretto, J., van Donselaar, E., Neri, A., D’Souza-Schorey, C., Chavrier, P., 1999. EFA6, a sec7 domain-containing exchange factor for ARF6, coordinates membrane recycling and actin cytoskeleton organization. EMBO J 18, 1480–1491.

Gasic, I., Mitchison, T.J., 2019. Autoregulation and repair in microtubule homeostasis. Curr Opin Cell Biol 56, 80–87.

Ghanta, K.S., Mello, C.C., 2020. Melting dsDNA Donor Molecules Greatly Improves Precision Genome Editing in *Caenorhabditis elegans*. Genetics 216, 643–650.

Ghosh, M., Lo, R., Ivic, I., Aguilera, B., Qendro, V., Devarakonda, C., Shapiro, L.H., 2019. CD13 tethers the IQGAP1-ARF6-EFA6 complex to the plasma membrane to promote ARF6 activation, beta1 integrin recycling, and cell migration. Sci Signal 12.

Honda, Y., Tsuchiya, K., Sumiyoshi, E., Haruta, N., Sugimoto, A., 2017. Tubulin isotype substitution revealed that isotype combination modulates microtubule dynamics in *C. elegans* embryos. J Cell Sci 130, 1652–1661.

Janke, C., Magiera, M.M., 2020. The tubulin code and its role in controlling microtubule properties and functions. Nat Rev Mol Cell Biol 21, 307–326.

Kamath, R.S., Fraser, A.G., Dong, Y., Poulin, G., Durbin, R., Gotta, M., Kanapin, A., Le Bot, N., Moreno, S., Sohrmann, M., Welchman, D.P., Zipperlen, P., Ahringer, J., 2003. Systematic functional analysis of the *Caenorhabditis elegans* genome using RNAi. Nature 421, 231–237.

Kanamarlapudi, V., 2014. Exchange factor EFA6R requires C-terminal targeting to the plasma membrane to promote cytoskeletal rearrangement through the activation of ADP-ribosylation factor 6 (ARF6). J Biol Chem 289, 33378–33390.

Kato, M., McKnight, S.L., 2018. A Solid-State Conceptualization of Information Transfer from Gene to Message to Protein. Annu Rev Biochem 87, 351–390.

King, M.R., Petry, S., 2020. Phase separation of TPX2 enhances and spatially coordinates microtubule nucleation. Nat Commun 11, 270.

Klein, S., Partisani, M., Franco, M., Luton, F., 2008. EFA6 facilitates the assembly of the tight junction by coordinating an Arf6-dependent and -independent pathway. J Biol Chem 283, 30129–30138.

Laan, L., Pavin, N., Husson, J., Romet-Lemonne, G., van Duijn, M., Lopez, M.P., Vale, R.D., Julicher, F., Reck-Peterson, S.L., Dogterom, M., 2012. Cortical dynein controls microtubule dynamics to generate pulling forces that position microtubule asters. Cell 148, 502–514.

Lawrence, E.J., Zanic, M., 2019. Rescuing microtubules from the brink of catastrophe: CLASPs lead the way. Curr Opin Cell Biol 56, 94–101.

Luton, F., Klein, S., Chauvin, J.P., Le Bivic, A., Bourgoin, S., Franco, M., Chardin, P., 2004. EFA6, exchange factor for ARF6, regulates the actin cytoskeleton and associated tight junction in response to E-cadherin engagement. Mol Biol Cell 15, 1134–1145.

Milanini, J., Fayad, R., Partisani, M., Lecine, P., Borg, J.P., Franco, M., Luton, F., 2018. EFA6 proteins regulate lumen formation through alpha-actinin 1. J Cell Sci 131.

Moreci, R.S., Lechler, T., 2021. KIF18B is a cell type-specific regulator of spindle orientation in the epidermis. Mol Biol Cell 32, ar29.

Morrissette, N.S., Mitra, A., Sept, D., Sibley, L.D., 2004. Dinitroanilines bind alpha-tubulin to disrupt microtubules. Mol Biol Cell 15, 1960–1968.

Nashchekin, D., Fernandes, A.R., St Johnston, D., 2016. Patronin/Shot Cortical Foci Assemble the Noncentrosomal Microtubule Array that Specifies the Drosophila Anterior-Posterior Axis. Dev Cell 38, 61–72.

Nishida, K., Tsuchiya, K., Obinata, H., Onodera, S., Honda, Y., Lai, Y.C., Haruta, N., Sugimoto, A., 2021. Expression Patterns and Levels of All Tubulin Isotypes Analyzed in GFP Knock-In *C. elegans* Strains. Cell Struct Funct 46, 51–64.

O’Rourke, S.M., Christensen, S.N., Bowerman, B., 2010. *Caenorhabditis elegans* EFA-6 limits microtubule growth at the cell cortex. Nat Cell Biol 12, 1235–1241.

O’Rourke, S.M., Dorfman, M.D., Carter, J.C., Bowerman, B., 2007. Dynein modifiers in C. elegans: light chains suppress conditional heavy chain mutants. PLoS Genet 3, e128.

Ohi, R., Strothman, C., Zanic, M., 2021. Impact of the ‘tubulin economy’ on the formation and function of the microtubule cytoskeleton. Curr Opin Cell Biol 68, 81–89.

Partisani, M., Baron, C.L., Ghossoub, R., Fayad, R., Pagnotta, S., Abelanet, S., Macia, E., Brau, F., Lacas-Gervais, S., Benmerah, A., Luton, F., Franco, M., 2021. EFA6A, an exchange factor for Arf6, regulates early steps in ciliogenesis. J Cell Sci 134.

Phillips, J.B., Lyczak, R., Ellis, G.C., Bowerman, B., 2004. Roles for two partially redundant alpha-tubulins during mitosis in early *Caenorhabditis elegans* embryos. Cell Motil Cytoskeleton 58, 112–126.

Qu, Y., Hahn, I., Lees, M., Parkin, J., Voelzmann, A., Dorey, K., Rathbone, A., Friel, C.T., Allan, V.J., Okenve-Ramos, P., Sanchez-Soriano, N., Prokop, A., 2019. Efa6 protects axons and regulates their growth and branching by inhibiting microtubule polymerisation at the cortex. Elife 8.

Quintin, S., Wang, S., Pontabry, J., Bender, A., Robin, F., Hyenne, V., Landmann, F., Gally, C., Oegema, K., Labouesse, M., 2016. Non-centrosomal epidermal microtubules act in parallel to LET-502/ROCK to promote *C. elegans* elongation. Development 143, 160–173.

Sakagami, H., 2008. The EFA6 family: guanine nucleotide exchange factors for ADP ribosylation factor 6 at neuronal synapses. Tohoku J Exp Med 214, 191–198.

Singh, D., Schmidt, N., Muller, F., Bange, T., Bird, A.W., 2021. Destabilization of Long Astral Microtubules via Cdk1-Dependent Removal of GTSE1 from Their Plus Ends Facilitates Prometaphase Spindle Orientation. Curr Biol 31, 766–781 e768.

Taffoni, C., Omi, S., Huber, C., Mailfert, S., Fallet, M., Rupprecht, J.F., Ewbank, J.J., Pujol, N., 2020. Microtubule plus-end dynamics link wound repair to the innate immune response. Elife 9.

Uversky, V.N., 2017. Intrinsically disordered proteins in overcrowded milieu: Membrane-less organelles, phase separation, and intrinsic disorder. Curr Opin Struct Biol 44, 18–30.

Wang, S., Wu, D., Quintin, S., Green, R.A., Cheerambathur, D.K., Ochoa, S.D., Desai, A., Oegema, K., 2015. NOCA-1 functions with gamma-tubulin and in parallel to Patronin to assemble non-centrosomal microtubule arrays in *C. elegans*. Elife 4, e08649.

White, J.G., Southgate, E., Thomson, J.N., Brenner, S., 1986. The structure of the nervous system of the nematode *Caenorhabditis elegans*. Philos Trans R Soc Lond B Biol Sci 314, 1–340.

Woodruff, J.B., Ferreira Gomes, B., Widlund, P.O., Mahamid, J., Honigmann, A., Hyman, A.A., 2017. The Centrosome Is a Selective Condensate that Nucleates Microtubules by Concentrating Tubulin. Cell 169, 1066–1077 e1010.

Yogev, S., Cooper, R., Fetter, R., Horowitz, M., Shen, K., 2016. Microtubule Organization Determines Axonal Transport Dynamics. Neuron 92, 449–460.

Zheng, Q., Schaefer, A.M., Nonet, M.L., 2011. Regulation of *C. elegans* presynaptic differentiation and neurite branching via a novel signaling pathway initiated by SAM-10. Development 138, 87–96.

Ziel, J.W., Hagedorn, E.J., Audhya, A., Sherwood, D.R., 2009. UNC-6 (netrin) orients the invasive membrane of the anchor cell in *C. elegans*. Nat Cell Biol 11, 183–189.

