## Supplemental Figures for "The microtubule regulator EFA-6 forms spatially restricted cortical foci dependent on its intrinsically disordered region and interactions with tubulins"

**Sandhu, Lyu et al. Supplemental Figures and Legends**

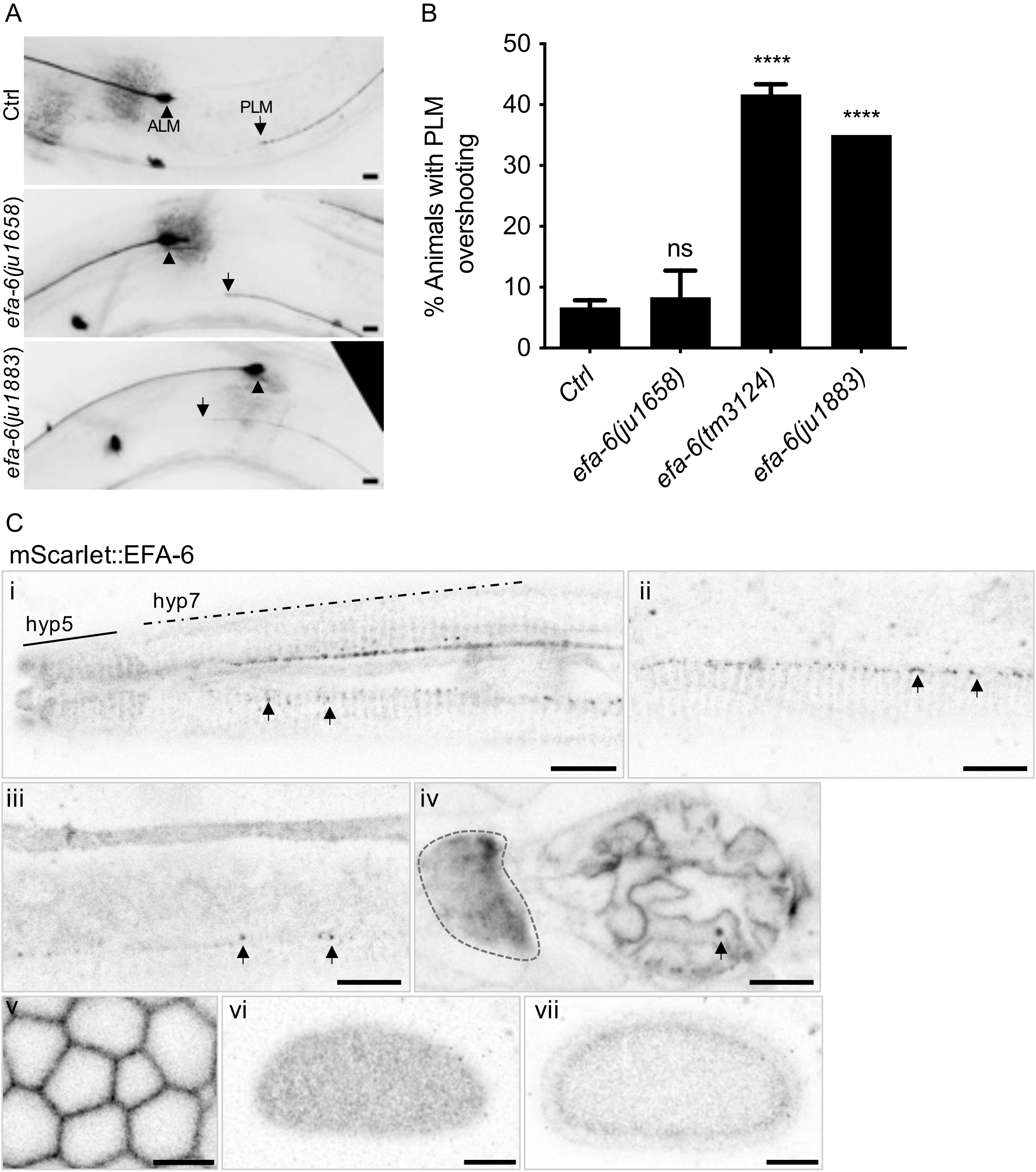


**Figure S1. Validation of EFA-6 fluorescent protein knock-ins**

(A) Representative touch neuron morphology images of *jsIs973*(P*mec-7*-mRFP) at mid-body region in *efa-6* knock-in and loss of function mutants. In control and GFP::EFA-6(*ju1658*) knock-in animals, PLM anterior neurites (arrows) terminated posterior to ALM soma (arrowheads). In *efa-6*(*ju1883*) null mutants and *efa-6(tm3124)* loss of function mutants, the PLM anterior neurite displayed overshooting i.e. termination anterior to ALM soma. Scale = 10 µm. (B) Quantitation of percentage of animals showing PLM overshooting. N = 45 per genotype, Statistics, Fisher Exact Test, ns, not significant; **** $P \leq 0.0001$. (C) mSc::EFA-6(*syb6998*) knock-in showed punctate (arrows) and diffuse pattern at the cellular cortex of epidermis (i-iii), nerve ring (iv, outlined area) and pharyngeal cells (iv, foci marked by arrow), germline (v), and single cell stage embryos (vi,vii). Longitudinal extent of hyp5 and hyp7 syncytia shown in (i) by solid and dashed lines respectively. Scale = 10 µm, except for C.v. = 5 µm.

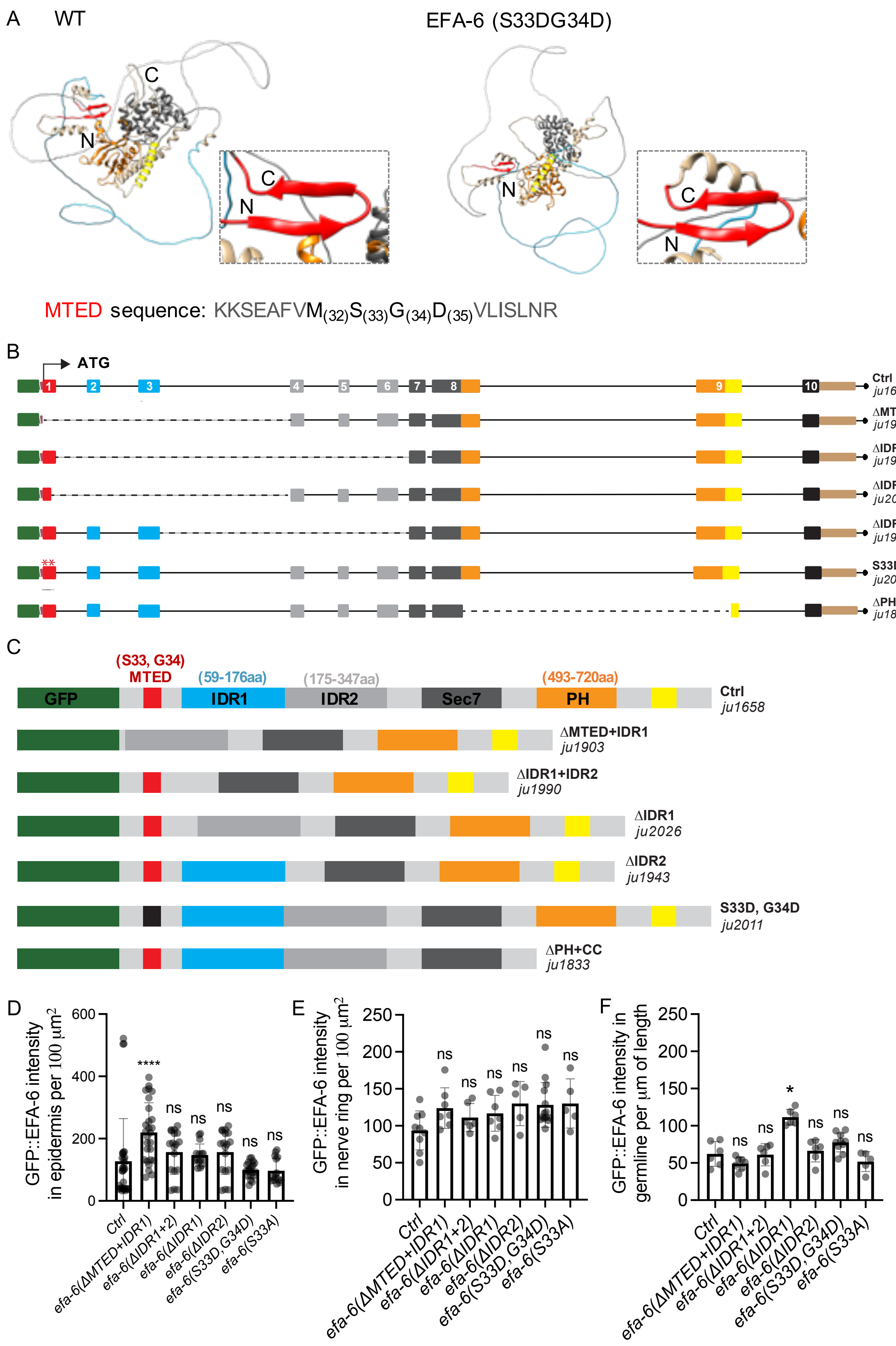
A
WT
EFA-6 (S33DG34D)
C
N
MTED sequence: KKSEAFVM(32)S(33)G(34)D(35)VLISLNR
B
ATG
Ctrl
ju1658
ΔMTED+IDR1
ju1903
ΔIDR1+IDR2
ju1990
ΔIDR1
ju2026
ΔIDR2
ju1943
S33D, G34D
ju2011
ΔPH+CC
ju1833
C
(S33, G34)
MTED
(59-176aa)
IDR1
(175-347aa)
IDR2
Sec7
(493-720aa)
PH
GFP
D
GFP::EFA-6 intensity in epidermis per 100 μm²
****
ns
E
GFP::EFA-6 intensity in nerve ring per 100 μm²
F
GFP::EFA-6 intensity in germline per μm of length
*
Ctrl
efa-6(ΔMTED+IDR1)
efa-6(ΔIDR1+2)
efa-6(ΔIDR1)
efa-6(ΔIDR2)
efa-6(S33D,G34D)
efa-6(S33A)

**Figure S2. Predicted EFA-6 protein structure and location of mutant alleles**

(A) EFA-6 structure predictions from Alphafold. The Sec7 and PH domains were predicted with high confidence whereas a structure was not predicted for the IDR (amino acids 51-377). The N-terminal Microtubule Elimination Domain (MTED, residues 25-42) was predicted to form a beta-sheet hairpin loop (shown in enlarged view on the right) containing a core of highly conserved residues: $M_{32}S_{33}G_{34}D_{35}$. Mutation of these residues (e.g. S33D, G34D) was predicted to not affect this loop. (B) *efa-6* genomic organization illustrating deletion or missense alleles generated by CRISPR/Cas9 engineering in the GFP::EFA-6(*ju1658*) background. Boxes represent exons, lines for introns, dashed lines mark deleted sequences, and asterisks mark editing within MTED. Colored lines above correspond to depicted domains of EFA-6. (C) Illustration of mutated GFP::EFA-6 proteins produced in the edited alleles, based on cDNA sequencing except ΔPH+CC. (D-E). Quantitation of GFP::EFA-6 intensity in anterior epidermis, nerve ring and germline. Statistics, Kruskal-Wallis test; ns, not significant, *, $P \leq 0.05$, **** $P \leq 0.0001$.

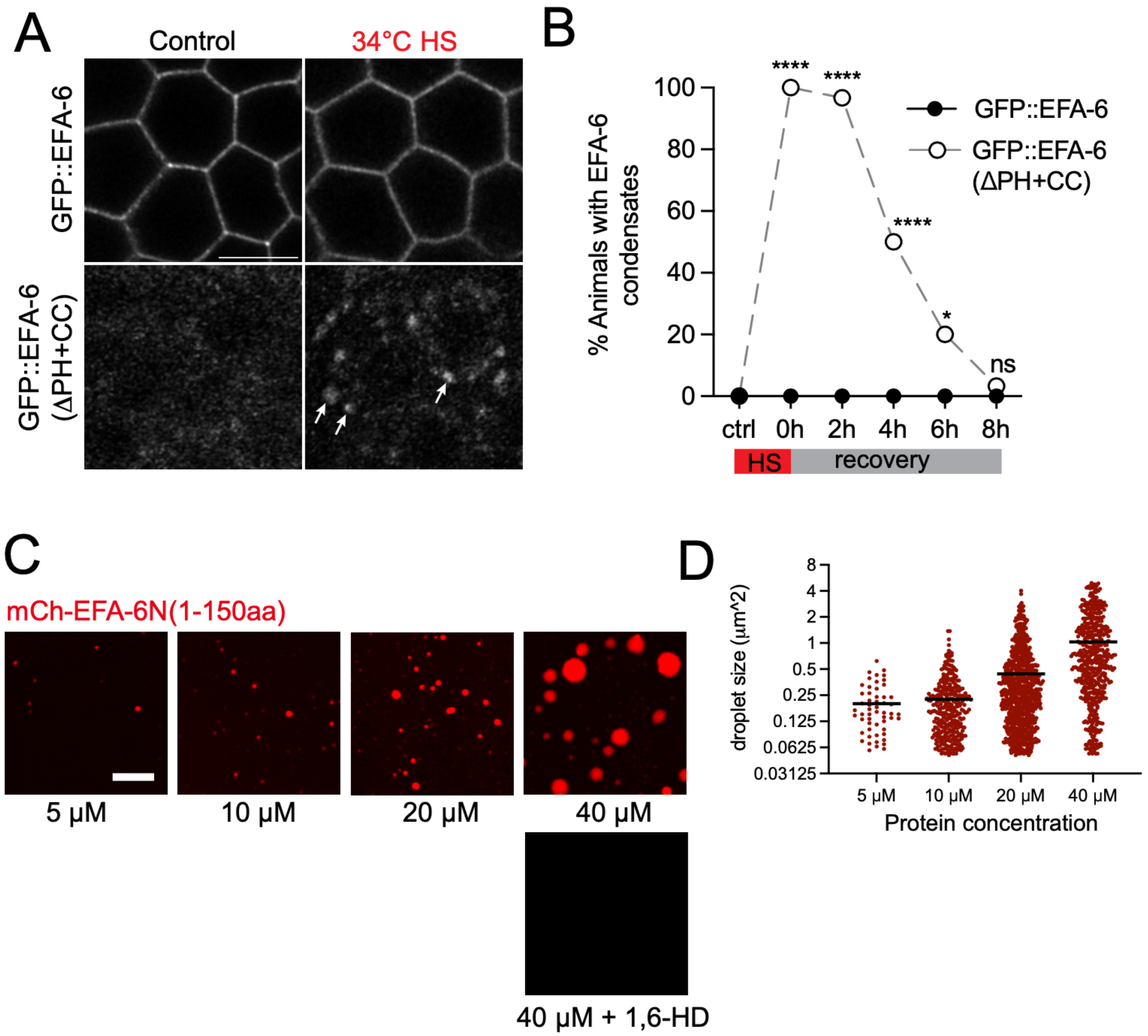


**Figure S3. The EFA-6 N-terminal domains can form punctate foci after heat stress *in vivo*, and purified EFA-6 N-terminal 150aa is sufficient to form condensates *in vitro***

(A) Shown are Airyscan images of GFP::EFA-6 and GFP::EFA-6(ΔPH+CC) in the syncytial germline. Arrows indicate GFP::EFA-6(ΔPH+CC) foci forming in germline after 2 h at 34°C. Wild type GFP::EFA-6 did not form condensates after 2 h at 34°C. Scale = 5 µm. (B) Quantitation of percentage of animals showing condensates at different time points of recovery at room temperature after 2 h heat shock (-2 to 0 h). Controls were grown at 20°C and imaged without heat shock. N = 30 animals per time point per genotype. Statistics, Fisher Exact Test, ns, not significant; **** $P \leq 0.0001$; * $P \leq 0.05$. (C, D) EFA-6N formed concentration-dependent condensates *in vitro* and the condensates were sensitive to 1,6-Hexanediol (1,6-HD). Representative images (C) and quantitation (D) for droplets of purified mCh-EFA-6N(1-150) at indicated protein concentrations in droplet formation buffer containing 20% PEG8000 (or droplet formation buffer containing 20% PEG8000 and 10% 1,6-Hexanediol). The presence of 1,6-Hexanediol inhibited droplet formation. Scale = 10 µm. Quantitation: n >3 independent trials.

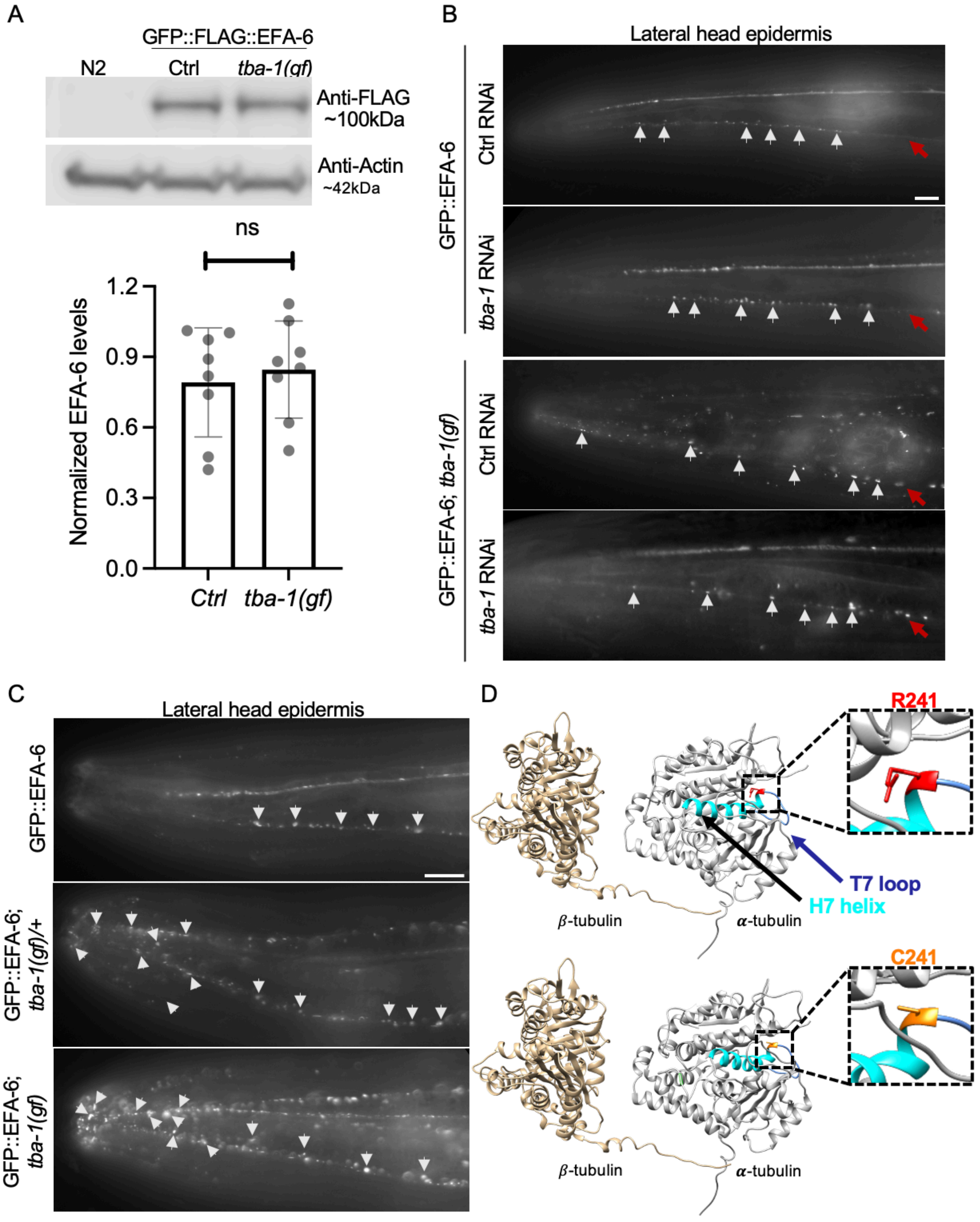


**Figure S4: TBA-1 α-tubulin predicted structure and dominant effect of R241C on EFA-6 condensates**

(A) EFA-6 protein levels were not affected in *tba-1(gf)*. Representative western blot of wild type (N2) negative control), GFP::EFA-6 (positive control), and GFP::EFA-6; *tba-1(gf)* mutants using anti-FLAG antibody with anti-actin loading control is shown above. Graph of quantitation of GFP::EFA-6 levels normalized to actin in GFP::EFA-6; *tba-1(wt)* and GFP::EFA-6; *tba-1(gf)* is shown below (n = 3 lysates; each dot represents one blot). Statistics: t test; ns, not significant. (B) GFP::EFA-6 localization in the lateral head epidermis in wild type and *tba-1(gf)* background, after control or *tba-1* RNAi. White arrows indicate GFP::EFA-6 condensates, red arrows indicate epidermal ridge margin. *tba-1* RNAi did not affect normal GFP::EFA-6 condensates but rescued the ectopic GFP::EFA-6 condensates observed in *tba-1(gf)*. Scale = 10 µm. (C) GFP::EFA-6 in the lateral head epidermis in wildtype, *tba-1(ju1869)*/+ heterozygous and homozygous background. *tba-1(ju1869)*/+ heterozygous animals and *tba-1(ju1869)* homozygotes showed similar ectopic GFP::EFA-6 condensates in head tip and along epidermal ridge margin. (D) α-tubulin TBA-1 structure modeled in UCSF Chimera showing position of R241 in the H7 helix. The TBA-1 GTP-binding pocket is at the other end of the H7 helix. R241 is in the interdimer α/β interface, potentially affecting dimer incorporation into MTs. R241C is not predicted to affect TBA-1 overall folding (WT shown above, TBA-1(gf) shown below).

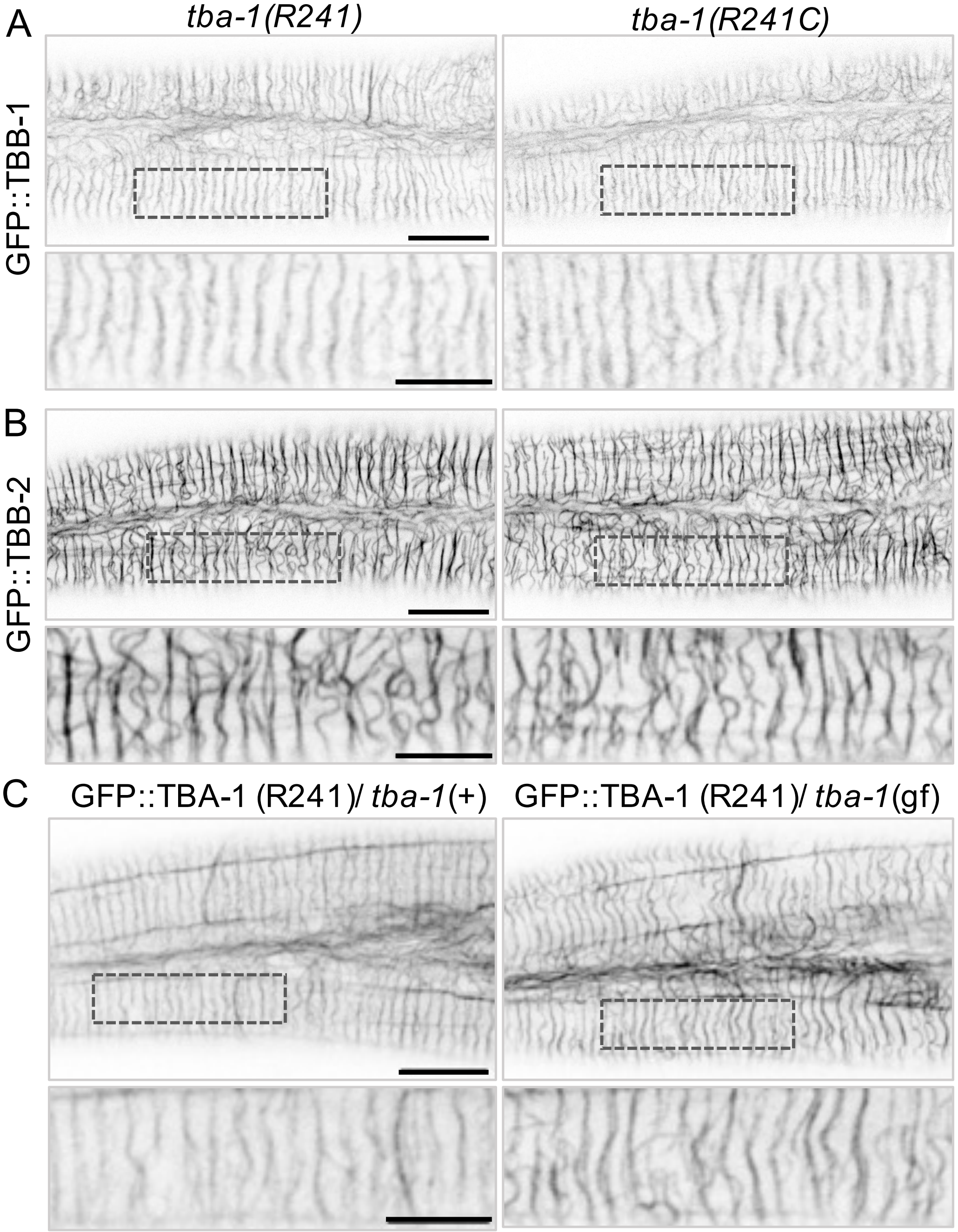


**Figure S5. *tba-1*(R241C) does not alter epidermal β-tubulin localization and shows reduced incorporation into MTs in trans-heterozygous *tba-1*(R241C) /GFP::TBA-1 animals**

(A, B) Shown are confocal single slice images of GFP::TBB-1 and GFP::TBB-2 knock-ins in control (*tba-1(R241)* and *tba-1(R241C, gf)* animals (1-day adult). Dashed boxes correspond to enlarged view below. Both markers displayed filamentous pattern in epidermis and were not strongly disrupted in *tba-1(gf)* mutants. (C) Confocal images of heterozygous GFP::TBA-1 (R241, wt) with *tba-1*(+ or *gf*). Dashed boxes correspond to enlarged images below. Epidermal GFP::TBA-1 filament intensity when heterozygous to untagged *tba-1(gf)* was increased compared to when heterozygous to wild type untagged *tba-1.* Scale = 10 µm for top images and 5 µm for enlarged images.

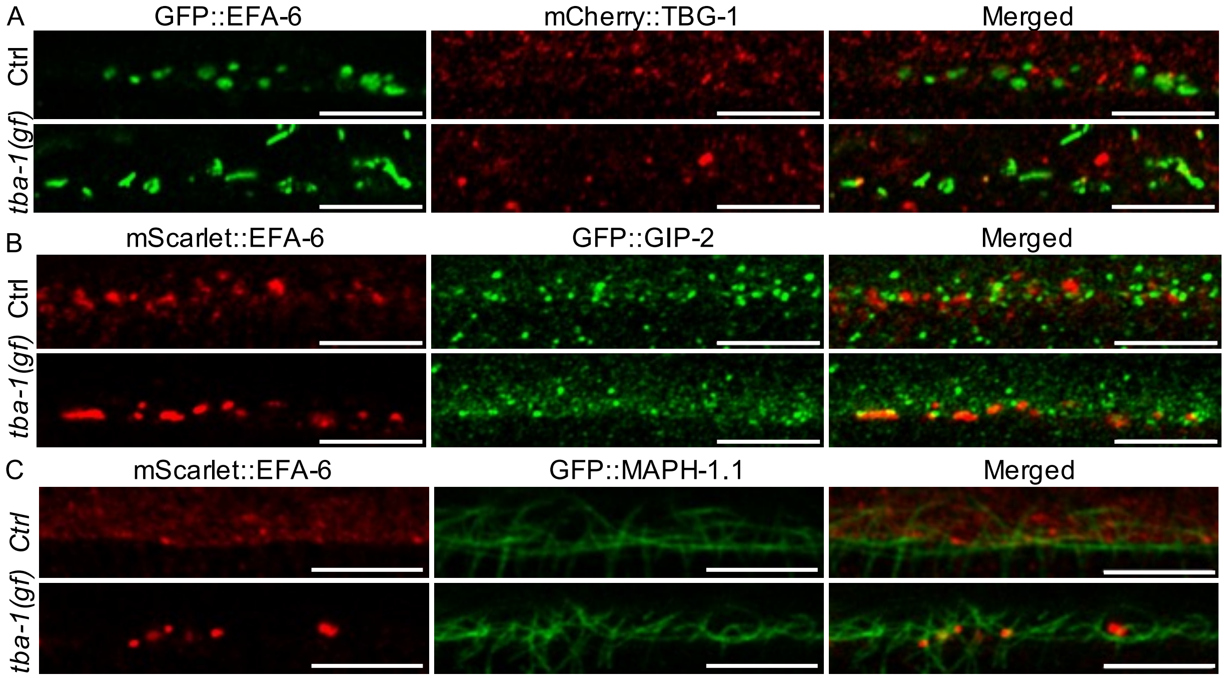


**Figure S6. GFP::EFA-6 condensates at epidermal ridge margin do not colocalize with MT markers in wild type or in *tba-1(gf)***

Representative airyscan single slice images of endogenously tagged EFA-6 at the epidermal ridge margin with mCherry::TBG-1(A), GFP::GIP-2 (B), or GFP::MAPH-1.1 (C) show no co-localization either in wild type or *tba-1(gf)* background. GFP::EFA-6 condensates in *tba-1(gf)* (panel A) are more variable in shape than in the wild type. Scale = 5 µm.

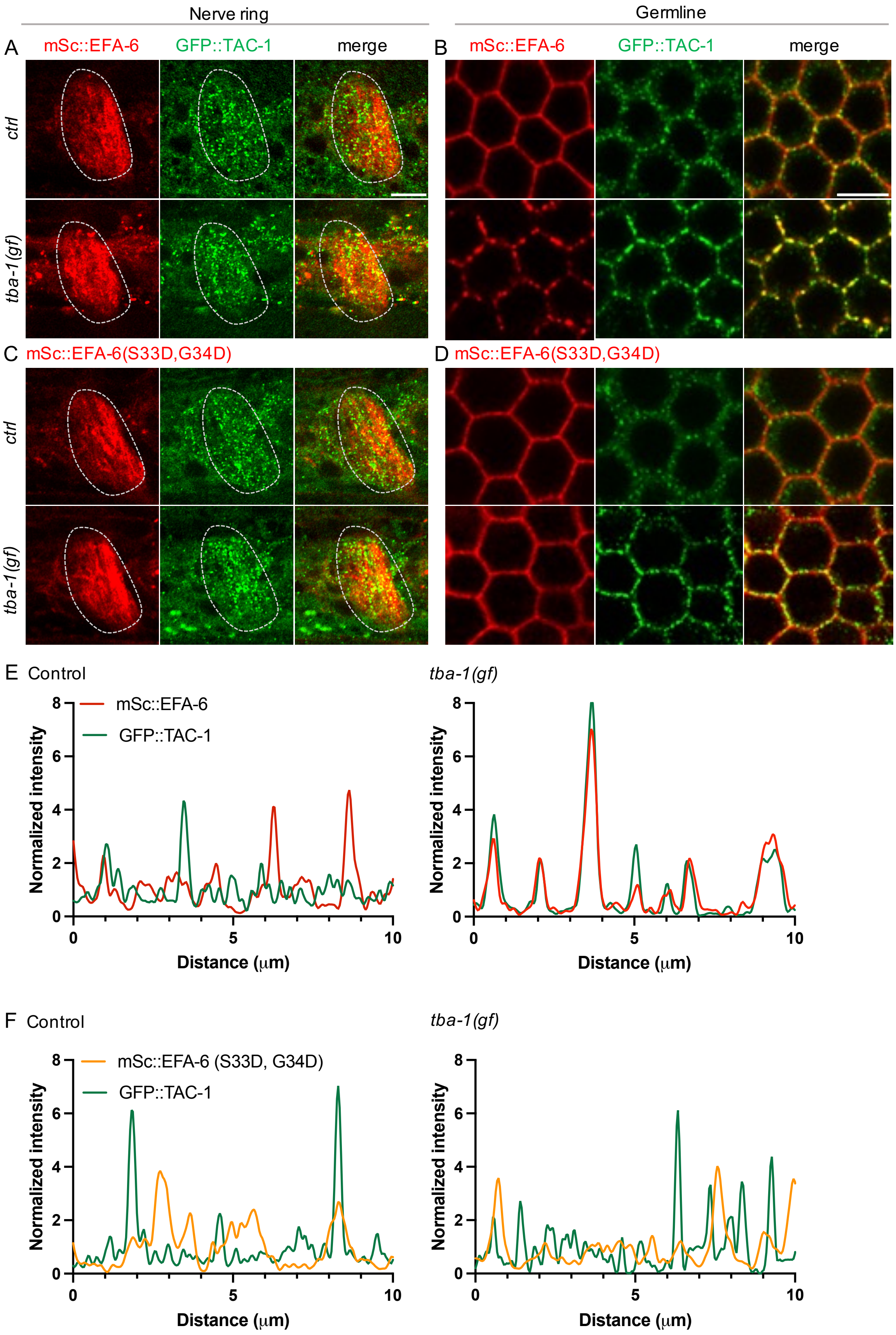
Nerve ring
Germline
A
mSc::EFA-6
GFP::TAC-1
merge
B
mSc::EFA-6
GFP::TAC-1
merge
ctrl
tba-1(gf)
C
mSc::EFA-6(S33D,G34D)
D
mSc::EFA-6(S33D,G34D)
ctrl
tba-1(gf)
E
Control
mSc::EFA-6
GFP::TAC-1
Normalized intensity
Distance (μm)
tba-1(gf)
Normalized intensity
Distance (μm)
F
Control
mSc::EFA-6 (S33D, G34D)
GFP::TAC-1
Normalized intensity
Distance (μm)
tba-1(gf)
Normalized intensity
Distance (μm)

**Figure S7. TAC-1 is recruited to *tba-1(gf)* induced mSc::EFA-6 condensates, dependent on the MTED of EFA-6**

(A, B) Shown are airyscan images of GFP::TAC-1 and mSc::EFA-6 in wild type and *tba-1(gf)* background, ROIs in nerve ring (A) and germline (B). GFP::TAC-1 formed puncta did not colocalize with mSc::EFA-6 in the wild type and colocalized in *tba-1(gf)*. (C, D) Shown are airyscan images of GFP::TAC-1 and mSc::EFA-6 (S33D,G34D) in wild type and *tba-1(gf)* background, ROIs in nerve ring (C) and germline (D). GFP::TAC-1 did not colocalize with mSc::EFA-6(S33D,G34D) in either background. Scale = 5 µm. (E, F) Representative line scans of mSc::EFA-6 (E) and mSc::EFA-6(S33D,G34D) (F) colocalization with GFP::TAC-1 in *tba-1(+)* control and *tba-1(gf)* mutants. Line scans of linear ROIs selected at the cortex of epidermal ridge margin (figures shown in Fig. 7C); intensity normalized to mean intensity of respective fluorescent markers.

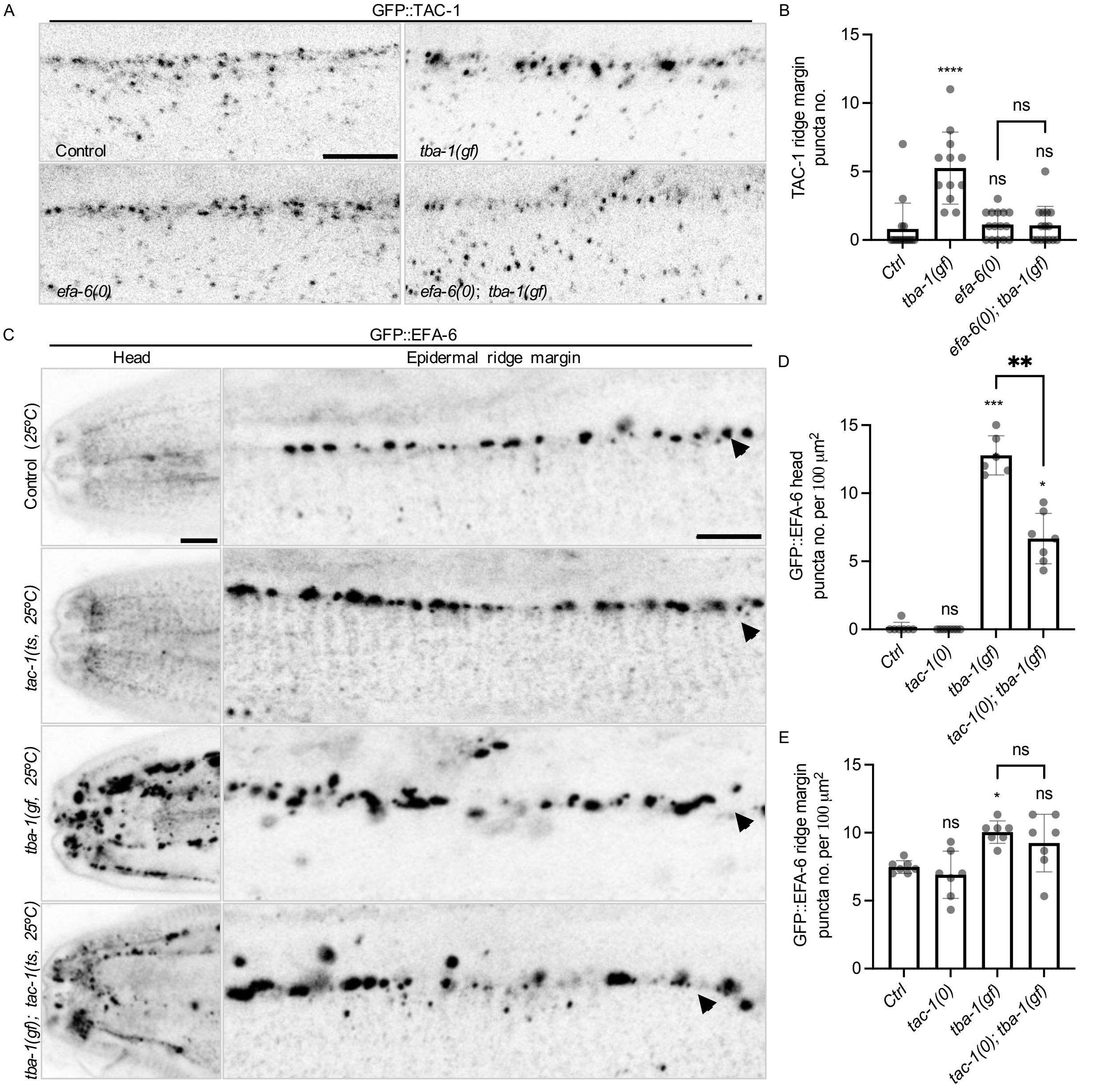


**Figure S8. Formation of large TAC-1 puncta in *tba-1(gf)* is suppressed by loss of function in *efa-6,* and loss of *tac-1* function partly reduces EFA-6 condensate formation**

(A) Airyscan single slice images of GFP::TAC-1 localization in epidermal ridge margin of *tba-1(+)* control, *tba-1(gf)* mutants, *efa-6(0)* null mutants, and *tba-1(gf);efa-6(0)* double mutants. Scale = 5 µm. (B) Quantitation of the density of large GFP::TAC-1 puncta (size larger than 0.13

μm$^2$) at the epidermal ridge margin, density was reduced in *tba-1(gf); efa-6(0)* compared to *tba-1(gf)*. Statistics: N = 5; Error bar, SEM; P value determined by Kruskal-Wallis test, ns (not significant), and **** P ≤ 0.0001. (C) Confocal images of GFP::EFA-6 localization in *tba-1(+)* control, *tac-1(or455ts)* loss of function mutants, *tba-1(gf)* mutants, and *tba-1(gf); tac-1(or455ts)* double mutants; all animals were imaged as day 1 adults after upshifting from 15°C to 25°C in late L3 stage. EFA-6 ectopic punctate condensate density was partially reduced in *tba-1(gf); tac-1(or455ts)* after temperature shift, relative to *tba-1(gf)*. For all images, scale = 5 μm; N = 6-7 animals per genotype. (D, E) Quantitation of GFP::EFA-6 condensate density at head and epidermal ridge margin. Loss of function in *tac-1* partially reduced GFP::EFA-6 condensate density per 100 μm$^2$ in head epidermis and in lateral epidermal ridge margins. Statistics: N = 5-10; Error bar, SEM; P value determined by Kruskal-Wallis test, ns (not significant) P > 0.05, * P ≤ 0.05, ** P ≤ 0.01, *** P ≤ 0.001, **** P ≤ 0.0001.
