## Supplemental Tables for "The microtubule regulator EFA-6 forms spatially restricted cortical foci dependent on its intrinsically disordered region and interactions with tubulins"

**Sandhu, Lyu, et al. Supplemental Tables 1-3**

**Supplemental Table 1. Key Resources**

| Reagent type (Species) or resource | Description | Source and/or reference | Catalog # RRID | Additional information |
|---|---|---|---|---|
| Antibody | Mouse Anti-actin monoclonal (Clone: C4) | MP Biomedicals | 0869100-CF | Western Blot, used at 1:10,000 |
| Antibody | Mouse Monoclonal Anti-alpha-tubulin (Clone: DM1A) | Sigma | T9026 | Western Blot, used at 1:5,000 |
| Antibody | Rabbit Anti-Beta-tubulin PolyAb | Proteintech | 10094-I-AP | Western Blot, used at 1:5,000 |
| Antibody | Rabbit Anti-FLAG | Sigma | F7425 | Western Blot, used at 1:5,000 |
| Western Blot Pre-prepared Gel | Bolt™ 4-12% Bis-Tris Plus | Invitrogen | NW04120BOX | Western Blot |
| DTT | NuPAGE | Invitrogen | NP0009 | Western Blot |
| LSD sample Buffer | NuPAGE | Invitrogen | NP0007 | Western Blot |
| Western Blot Running Buffer | Bolt™ MES SDS Running Buffer | Invitrogen | B0002 | Western Blot |
| Western Blot Transfer Buffer | Pierce™ Western Blot Transfer | Thermo Scientific | 35040 | Western Blot |
| Protein quantification kit | Pierce™ BCA Protein Assay Kit | Thermo Scientific | 23227 | Western Blot |
| Detergent | Tergitol™ Solution | Sigma | NP40S | Protein extraction buffer |
| Protease inhibitors | cOmplete™ Protease Inhibitor cocktail | Roche | 11873580001 | Protein extraction buffer |
| Phosphatase inhibitors | PhosSTOP EASYpack | Roche | 04 906 837 001 | Protein extraction buffer |
| Chemical | Sodium Molybdate dihydrate | Sigma | M1003 | Protein extraction buffer |
| Chemical | Sodium Fluoride | Sigma | S7920 | Protein extraction buffer |

| | | | | |
|---|---|---|---|---|
| Chemical | Phenylmethylsulfonyl fluoride | Sigma | P7626 | Protein extraction buffer |
| Chemical | Ni-NTA Resin | Thermo Fisher | 88221 | Recombinant protein purification |
| Chemical | PEG-8000 | Sigma | P2139 | Droplets formation buffer |
| Chemical | 1, 6 - Hexanediol | Sigma | 240117 | Droplets formation buffer |
| Genetic reagent (*C. elegans*) | | CGC | N2 | Wild type |
| Genetic reagent (*C. elegans*) | *gfp::efa-6(ju1658)* | This work | CZ27027 | KI used as reference control for all the figures |
| Genetic reagent (*C. elegans*) | *mSc::efa-6(syb6998)* | This work | CZ29957 | KI used as reference control for all the figures |
| Genetic reagent *(C. elegans)* | *mSc::efa-6(syb6998); lam-1::GFP(qyIs8)** | *(Ziel et al., 2009); This work | CZ30010 | Figure 1 |
| Genetic reagent (*C. elegans*) | *unc-119(ed3)*; *tbg-1::mCherry(ltSi62)** *gfp::efa-6(ju1658)* | *(Wang et al., 2015); This work | CZ29604 | Figure 1, S6 |
| Genetic reagent (*C. elegans*) | *gip-2::GFP(lt19)**; efa-6(syb6998)* | *(Wang et al., 2017); This work | CZ30170 | Figure 1, S6 |
| Genetic reagent (*C. elegans*) | *ebp-2::GFP(wow47)**; mSc::efa-6(syb6998)* | *(Sallee et al., 2018) | CZ30475 | Figure 1 |
| Genetic reagent (*C. elegans*) | P*mec-7*-mRFP + CB*unc-119* (*jsIs973*) | (Zheng et al., 2011) | NM3336 | Figure S1 |
| Genetic reagent (*C. elegans*) | P*mec-7*-mRFP(*jsIs973*); *efa-6(ju1658)* | This work | CZ26544 | Figure S1 |
| Genetic reagent (*C. elegans*) | P*mec-7*-mRFP(*jsIs973*); *efa-6(tm3124)* | This work | CZ28608 | Figure S1 |
| Genetic reagent (*C. elegans*) | P*mec-7*-mRFP(*jsIs973*); *efa-6(ju1883ju1658)* | This work | CZ30221 | Figure S1 |
| Genetic reagent (*C. elegans*) | *efa-6(ju1903ju1658)* | This work | CZ29095 | Figure 2, Table 1 |
| Genetic reagent (*C. elegans*) | P*mec-7*-mRFP(*jsIs973*); *efa-6(ju1990ju1658)* | This work | CZ29798 | Figure 2, Table 1 |

| | | | | |
|---|---|---|---|---|
| Genetic reagent (*C. elegans*) | *efa-6(ju2026ju1658)* | This work | CZ30298 | Figure 2 |
| Genetic reagent (*C. elegans*) | *efa-6(ju1943ju1658)* | This work | CZ29509 | Figure 2 |
| Genetic reagent (*C. elegans*) | *efa-6(ju2011ju1658)* | This work | CZ29963 | Figure 2, Table 1 |
| Genetic reagent (*C. elegans*) | *efa-6(ju1974ju1658)* | This work | CZ29673 | Figure 2 |
| Genetic reagent (*C. elegans*) | *efa-6(ju1833ju1658)* | This work | CZ28317 | Figure S3 |
| Genetic reagent (*C. elegans*) | *tba-1(ju1761); gfp::efa-6(ju1658)* | This work | CZ27301 | Original isolate of *tba-1(ju1761)* |
| Genetic reagent (*C. elegans*) | *tba-1(ju1761); gfp::efa-6(ju1658)* | This work | CZ27771 | Outcrossed *tba-1(ju1761)* |
| Genetic reagent (*C. elegans*) | *tba-1(ju1869); gfp::efa-6(ju1658)* | This work | CZ28692 | *tba-1(1869)* made by CRISPR-editing |
| Genetic reagent (*C. elegans*) | *tba-1(ok1135); gfp::efa-6(ju1658)* | This work | CZ28682 | Figure 3, 7 |
| Genetic reagent (*C. elegans*) | *efa-6(ju1883ju1658)* | This work | CZ30053 | Table 1 |
| Genetic reagent (*C. elegans*) | *tba-1(ju1869); efa-6(ju1883ju1658)* | This work | CZ30305 | Table 1 |
| Genetic reagent (*C. elegans*) | *tba-1(ju1872); gfp::efa-6(ju1658)* | This work | CZ28911 | Table 2 |
| Genetic reagent (*C. elegans*) | *tba-1(ju1876)/ [bli-4(e937) let-?(q782) qIs48](hT2); gfp::efa-6(ju1658)* | This work | CZ28918 | Table 2 |
| Genetic reagent (*C. elegans*) | *tba-1(ju1871)/ [bli-4(e937) let-?(q782) qIs48](hT2); gfp::efa-6(ju1658)* | This work | CZ28920 | Table 2 |
| Genetic reagent (*C. elegans*) | *tba-1(ju1929); gfp::efa-6(ju1658)* | This work | CZ29489 | Table 2 |
| Genetic reagent (*C. elegans*) | *tba-2(ju1875)/ In(ile-1 Y18D10A.2 In(dnj-27 dkf-1)) (tmC27)*; gfp::efa-6(ju1658)* | *(Dejima et al., 2018); This work | CZ28915 | Table 2 |
| Genetic reagent (*C. elegans*) | *tbb-1(gk207); gfp::efa-6(ju1658)* | This work | CZ30487 | Figure 4 |
| Genetic reagent (*C. elegans*) | *tba-1(ju1869); tbb-1(gk207); gfp::efa-6(ju1658)* | This work | CZ29512 | Figure 4 |
| Genetic reagent (*C. elegans*) | *tbb-2(gk129); gfp::efa-6(ju1658)* | This work | CZ29510 | Figure 4 |

| | | | | |
|---|---|---|---|---|
| Genetic reagent (*C. elegans*) | *tba-1(ju1869); tbb-2(gk129); gfp::efa-6(ju1658)* | This work | CZ29511 | Figure 4 |
| Genetic reagent (*C. elegans*) | *evl-20(ar103)*/ dpy-10(e128) unc-4(e120); gfp::efa-6(ju1658)* | *(Antoshechkin and Han, 2002); This work | CZ29146 | Figure 4 |
| Genetic reagent (*C. elegans*) | *tba-1(ju1869); evl-20(ar103) / dpy-10(e128) unc-4(e120); gfp::efa-6(ju1658)* | This work | CZ29709 | Figure 4 |
| Genetic reagent (*C. elegans*) | GFP::TEV::3xFLAG::*tba-1(tj44)* | (Honda et al., 2017) | SA1067 | Figure 5 |
| Genetic reagent (*C. elegans*) | GFP::TEV::3xFLAG::*tba-1(ju1868 tj44)* | This work | CZ28693 | Figure 5 |
| Genetic reagent (*C. elegans*) | *GFP::maph-1.1(mib12)* | (Waaijers et al., 2016) | BOX188 | Figure 5 |
| Genetic reagent (*C. elegans*) | *tba-1(ju1869); GFP::maph-1.1(mib12* | This work | CZ28814 | Figure 5 |
| Genetic reagent (*C. elegans*) | *GFP::tbb-1(tj30)* | (Honda et al., 2017) | SA884 | Figure S5 |
| Genetic reagent (*C. elegans*) | *tba-1(ju1869); GFP::tbb-1(tj30)* | This work | CZ29502 | Figure S5 |
| Genetic reagent (*C. elegans*) | *GFP::tbb-2(tj26)* | (Honda et al., 2017) | SA874 | Figure S5 |
| Genetic reagent (*C. elegans*) | *tba-1(ju1869); GFP::tbb-2(tj26)* | This work | CZ29501 | Figure S5 |
| Genetic reagent (*C. elegans*) | *GFP::maph-1.1(mib12); mSc::efa-6(syb6998)* | This work | CZ29964 | Figure S6 |
| Genetic reagent (*C. elegans*) | *tba-1(ju1869); GFP::maph-1.1(mib12); mSc::efa-6(syb6998)* | This work | CZ30082 | Figure S6 |
| Genetic reagent (*C. elegans*) | *tba-1(ju1869)*; *unc-119(ed3); tbg-1::mCherry(ltsi62)*; *gfp::efa-6(ju1658)* | This work | CZ29518 | Figure S6 |
| Genetic reagent (*C. elegans*) | *tba-1(ju1869); gip-2::GFP(lt19); efa-6(syb6998)* | This work | CZ29954 | Figure S6 |
| Genetic reagent (*C. elegans*) | *tba-1(ju1869); efa-6(ju1903ju1658)* | This work | CZ29241 | Figure 6, Table 1 |
| Genetic reagent (*C. elegans*) | *tba-1(ju1869); efa-6(ju1990ju1658)* | This work | CZ30009 | Figure 6, Table 1 |
| Genetic reagent (*C. elegans*) | *tba-1(ju1869);* $P_{mec-7}$-mRFP(*jsIs973*); *efa-6(ju2026ju1658)* | This work | CZ30299 | Figure 6 |
| Genetic reagent (*C. elegans*) | *tba-1(ju1869); efa-6(ju1943ju1658)* | This work | CZ29514 | Figure 6 |

| | | | | |
|---|---|---|---|---|
| Genetic reagent (*C. elegans*) | *tba-1(ju1869); efa-6(ju2011ju1658)* | This work | CZ30331 | Figure 6, Table 1 |
| Genetic reagent (*C. elegans*) | *gfp::tac-1(or1955)*; efa-6(syb6998)* | *(Chuang et al., 2020); This work | CZ30169 | Figure 7, S7 |
| Genetic reagent (*C. elegans*) | *tba-1(ju1869); gfp::tac-1(or1955); efa-6(syb6998)* | This work | CZ29955 | Figure 7, S7 |
| Genetic reagent (*C. elegans*) | *GFP::tac-1(or1955); efa-6(ju2028syb6998)* | This work | CZ30311 | Figure 7, S7 |
| Genetic reagent (*C. elegans*) | *tba-1(ju1869); gfp::tac-1(or1955); efa-6(ju2028syb6998)* | This work | CZ30312 | Figure 7, S7 |
| Genetic reagent (*C. elegans*) | *GFP::tac-1(or1955); efa-6(ju1883ju1658)* | This work | CZ30249 | Figure S8 |
| Genetic reagent (*C. elegans*) | *tba-1(ju1869); GFP::tac-1(or1955); efa-6(ju1883ju1658)* | This work | CZ30251 | Figure S8 |
| Genetic reagent (*C. elegans*) | *tac-1(or455)*; gfp::efa-6(ju1658)* | *(Bellanger et al., 2007); This work | CZ30247 | Figure S8 |
| Genetic reagent (*C. elegans*) | *tba-1(ju1869); tac-1(or455); gfp::efa-6(ju1658)* | This work | CZ30248 | Figure S8 |
| Oligonucleotide | 5'-CGTTTTCAGAGTGATGGCGAGTTTTAGAGCTAGAAATAGCAAG-3' | This work | YJ12362 | used to generate pCZ986 |
| Oligonucleotide | 5'-TCGCCATCACTCTGAAAACGCAAGACATCTCGCAATAGGAGG-3' | This work | YJ12363 | used to generate pCZ986 |
| Oligonucleotide | 5'-ACGTTGTAAAACGACGGCCAGTCGCCGGCAAAACCCTCCTTACCGCCTAATCAC-3' | This work | YJ12364 | used to generate pCZ996 |
| Oligonucleotide | 5'-TCCAGTGAACAATTCTTCTCCTTTACTCATCACTCTGAAAACGAGATGATTGTAG-3' | This work | YJ12365 | used to generate pCZ996 |
| Oligonucleotide | 5'-CGTGATTACAAGGATGACGATGACAAGAGAATGGCGAAAGTCGCGTCGTC-3' | This work | YJ12369 | used to generate pCZ996 |

| | | | | |
|---|---|---|---|---|
| Oligonucleotide | 5’-TCACACAGGAAACAGCTATGACCATGTTATACCACGTCTTTCGATTTGGCTTG-3’ | This work | YJ12370 | used to generate pCZ996 |
| Plasmid | pDD282: GFP^SEC^3xFlag vector with ccdB markers for cloning homology arms | (Dickinson et al., 2015) | Addgene #66823 | Used to generate pCZ996 |
| Plasmid | pDD162: Peft-3-Cas9 + Empty sgRNA | (Dickinson et al., 2015) | Addgene #47549 | Used to generate pCZ986 |
| Plasmid | pCZ996 | This work | | Contain homology arms of *efa-6* used in generating *ju1658* |
| Plasmid | pCZ986 | This work | | Contain targeting sgRNA of *efa-6* and used in generating *ju1658* |
| Plasmid | pCHN261 pET-His-mCh-EFA-6N | This work | | Used to generate recombinant protein mCh-EFA-6N |
| Plasmid | pCHN313 pET-His-GFP-TAC-1 | This work | | Used to generate recombinant protein GFP-TAC-1 |

**Supplemental Table 2: Allele information and detection methods**

| Allele name | Genotyping primers, PCR products and detection methods | Allele information and predicted protein |
|---|---|---|
| *efa-6(ju1658)* | Forward: ctcacctccatcacgtgttt (YJ12371)<br>Reverse: tgaggcctcaaatcgcagagc (YJ12372)<br>WT: 1542 bp<br>Knock-in (KI): 507 bp | GFP insertion in-frame to ATG of endogenous EFA-6* |
| *efa-6(syb6998)* | Forward: gcgttccgagggacgtcact (SD21116)<br>Reverse: tgaggcctcaaatcgcagagc (YJ12372)<br>WT: no band<br>KI: 429 bp | mScarlet insertion in-frame to ATG of endogenous EFA-6* |
| *efa-6(ju1903)* | Forward: caaacgagaagcgtgaccac (SD21117)<br>Reverse: gtggtgaaagtgtggatatttg (SD21118)<br>WT: 3126 bp<br>MUT: 343 bp | Deletion of nucleotide (nt) 1-2819 of *efa-6* genomic DNA*, corresponding to N-terminal ATG to E188, designated ΔMTED+IDR1. Sequencing of RT-PCR products verified the deletion and in-frame fusion of GFP with the remaining 629 aa of EFA-6. |
| | Forward: caaacgagaagcgtgaccac (SD21117)<br>Reverse: tgaggcctcaaatcgcagagc (YJ12372)<br>WT:625 bp<br>MUT: no band | |
| *efa-6(ju1943)* | Forward: atactcctccagtggctcctcgac (SD20090)<br>Reverse: atccgtttagaggttgtttagcactc (SD20088)<br>WT: 1760 bp<br>MUT: 610 bp | Deletion of nt 2797-3946 of genomic DNA, corresponding to deletion of Glu175 to Thr347, designated ΔIDR2. Sequencing of RT-PCR products verified the deletion and in-frame fusion of GFP with remaining 645 aa of EFA-6. |
| | Forward: ccgtcgaaacattcagtggt (YJ6254)<br>Reverse: gttgtggctgccgtcttgat (YJ6253)<br>WT:1021bp<br>MUT: no band | |
| *efa-6(ju2026)* | Forward: gaaataatgctctgcgatttgag (SD20611)<br>Reverse: ggttaaaaatcaatggaaaatcg (SD20610)<br>WT: 1113 bp<br>MUT: 450bp | Deletion of nt 2958-3731 of genomic DNA, corresponding to N-terminal P59 to F176, designated ΔIDR1. Sequencing of RT-PCR products verified the deletion and GFP in-frame fused to the remaining 699 aa. |
| *efa-6(ju1990)* | Forward: gaaataatgctctgcgatttgag (SD20611)<br>Reverse: atactcctccagtggctcctcgac (SD20090)<br>WT: 3764 bp<br>MUT: 360 bp | Deletion of nt 547-3945 of genomic DNA, corresponding to N-terminal G59 to T342 designated ΔIDR1+2. Sequencing of RT-PCR products verified the deletion and GFP in-frame fused to the remaining 534 aa |
| | Forward: gaaataatgctctgcgatttgag (SD20611) | |

| | | |
|---|---|---|
| | Reverse: ggttaaaaatcaatggaaaatcg (SD20610)<br>WT: 1113 bp<br>MUT: no band | |
| *efa-6(ju1974)* | Forward: caaacgagaagcgtgaccac (SD21117)<br>Reverse: tgaggcctcaaatcgcagagc (YJ12372)<br>Digest the 625 bp PCR product with TaqαI, WT product is cut into 389 + 236 bp, MUT not cut. | Editing of two nucleotides, t97g and c99t, resulting in S33A in MTED of EFA-6** |
| *efa-6(ju2011)* | Forward: caaacgagaagcgtgaccac (SD21117)<br>Reverse: tgaggcctcaaatcgcagagc (YJ12372)<br>Digest the 625 bp PCR product with TaqαI, WT product is cut into 389 + 236 bp, MUT not cut. | Editing of four nucleotides: t97g, c98a, c99, g101a, resulting in S33D and G34D in MTED of EFA-6** |
| *efa-6(ju2028)* | Forward: gcgttccgagggacgtcact (SD21116)<br>Reverse: tgaggcctcaaatcgcagagc (YJ12372)<br>Digest the 429bp PCR product with TaqαI, WT product is cut into 193 + 236 bp, MUT not cut. | Editing of four nucleotides: t97g, c98a, c99, g101a, resulting in S33D and G34D in MTED of EFA-6** |
| *efa-6(ju1833)* | Forward: gctcgaaatctgtatgagttgaaga(SD21128)<br>Reverse: ttgattctattgcgagaatatcttg(SD21129)<br>WT: 739 bp<br>MUT: no band<br><br>Forward: gctcgaaatctgtatgagttgaaga(SD21128)<br>Reverse: ctcgaaactcaaaagcacccat (SD21127)<br>WT: no band<br>MUT: 436 bp | Deletion of nt 4573-7697 of genomic DNA, corresponding to N-terminal H493 to H720, designated (ΔPH+CC); verified by Sanger sequencing. |
| *efa-6(ju1883)* | Forward: caaacgagaagcgtgaccac (SD21117)<br>Reverse: gtggtgaaagtgtggatatttg (SD21118)<br>WT: 513 bp<br>MUT: 82 bp<br><br>Forward: ttcagggagccggatctgat(YJ12370)<br>Reverse: gtggtgaaagtgtggatatttg (SD21118)<br>WT: no band<br>MUT: 159 bp | Deletion of nt 1-2799 of genomic DNA), corresponding to N-terminal ATG1 to E188, and resulting in out of frame followed by a stop codon, designated as null (0). RT-PCR sequencing verified GFP in-frame with 35 aa unrelated to EFA-6, however, no additional ATG would initiate the production of remaining C-terminal of EFA-6 protein. |
| *efa-6(tm3124)* | Forward: ccgtcgaaacattcagtggt (YJ6254)<br>Reverse: gttgtggctgccgtcttgat (YJ6253)<br>WT: 1021 bp<br>MUT: 450 bp | Deletion of nt 3913-4484 of genomic DNA, leading to out-of-frame after aa 345; RT-PCR sequencing suggests |

| | | |
|---|---|---|
| | | production N-terminal 344 aa may be present. |
| *tba-1(ju1869)* | Forward: gacatctcctctgtacaagag (SD20485)<br>Reverse: gcttccctccgtttcgatg (SD20930)<br>WT: 297 bp<br>MUT: no band | Editing of single nucleotide: c1062t, resulting in R241C**. |
| | Forward: gacatctcctctgtacaagag (SD20485)<br>Reverse: tctctttgttttgacggtgc (SD20929)<br>WT: no band<br>MUT: 297 bp | |
| *tba-1(ok1135)* | Forward: ctgtctcgagcatggaatccag (YJ11190)<br>Reverse: cgacctcttcgtagtccttttcg (YJ11191)<br>WT: 1324 bp<br>MUT: 288 bp | 1036 bp deletion of genomic DNA, nt 231-1267, resulting in premature stop codon. |
| *tba-1(ju1876)* | Forward: ccaagttacaccaacctgaac (SD21119)<br>Reverse: cagagatcagtggagtgtac (SD21120)<br>Digest the 173 bp PCR product with MnlI, WT product is cut into 100 + 73bp, MUT not cut. | Editing of three nucleotides: c1062a; g1063a, t1064g, resulting in R241K**. |
| *tba-1(ju1872)* | Forward: ccaagttacaccaacctgaac (SD21119)<br>Reverse: cagagatcagtggagtgtac (SD21120)<br>Digest the 173 bp PCR product with MnlI, WT product is cut into 100 + 73bp, MUT not cut | Editing of two nucleotides: c1062g, g1063c, resulting in R241A**. |
| *tba-1(ju1871)* | Forward: ccaagttacaccaacctgaac (SD21119)<br>Reverse: cagagatcagtggagtgtac (SD21120)<br>Digest the 173 bp PCR product with MnlI, WT product is cut into 100 bp+73bp, MUT isn't cut. | Editing of three nucleotides: c1062g, g1063a, t1064g, resulting in R241E**. |
| *tba-2(ju1875)* | Forward: caagctacaccaacctcaac (SD21121)<br>Reverse: agcagagatgagtggagtgt (SD21122)<br>Digest the 220 bp PCR product with TaqαI, WT product is cut into 117 + 103bp, MUT not cut. | Editing of two nucleotides: a771t, a773t, resulting in R241C**. |
| *GFP::tac-1(or1955)* | GFP signal visible under compound microscope | GFP insertion at the N-terminus of endogenous TAC-1 (Chuang et al., Biol. Open, 2020) |
| *gip-2::GFP::loxP::cb-unc-119(+)::loxP(lt19)* | GFP signal visible under compound microscope | Single-copy transgene with GFP fused at the C-terminus of GIP-2 (Wang et al., 2017) |
| *tbg-1::mCherry (ltSi62)* | mCherry signal visible under compound microscope | Single-copy transgene with mCherry fused at the C- |

| | | |
|---|---|---|
| | | terminus of TBG-1 (Wang et al., 2015) |
| *GFP::maph-1.1(mib12)* | GFP signal visible under dissection microscope | GFP insertion at the N-terminus of endogenous MAPH-1.1 (Waaijers et al., 2016) |
| *ebp-2::GFP::3xFLAG (wow47)* | GFP signal visible under compound microscope | GFP insertion at the C-terminus of endogenous EBP-2 (Sallee et al., 2018) |
| *GFP::TEV::3xFLAG::tba-1(tj44)* | GFP signal visible under dissection microscope | GFP insertion at the ATG of TBA-1 (Honda et al., J. Cell Sci., 2017) |
| *GFP::TEV::3xFLAG::tba-1(tj44) I tba-1(ju1868)* | Forward: gacatctcctctgtacaagag (SD20485)<br>Reverse: gcttccctccgtttcgatg (SD20930)<br>WT: 297 bp<br>MUT: no band | Editing of single nucleotide, c1062t, in *tj44*, resulting in GFP tagged TBA-1(R241C). |
| | Forward: gacatctcctctgtacaagag (SD20485)<br>Reverse: tctctttgttttgacggtgc (SD20929)<br>WT: no band<br>MUT: 297 bp | |
| *GFP::tbb-2(tj26)* | GFP signal visible under dissection scope | GFP insertion at the N-terminus of endogenous TBB-2 (Honda et al., 2017) |
| *GFP::tbb-1(tj30)* | GFP signal visible under dissection scope | GFP insertion at the N-terminus of endogenous TBB-1 (Honda et al., 2017) |
| *evl-20(ar103)* | Forward: gaattatttcgaatccacagatgctttaattt (SD21123)<br>Reverse: ttaatcaagtatgaataatcgacttcca (SD21124)<br>PCR product of 566 bp and sequencing | Contains a single nucleotide alteration, c468t, resulting in Q140 to TAG (Antoshechkin and Han, 2002) |
| *tbb-2(gk129)* | Forward: ccttgaagacaatcgcatccttcggc (YJ12613)<br>Reverse: gagcaaatagaaaggtacttgcgctg (YJ12614)<br>WT: 870 bp<br>MUT: 120 bp | Deletion of nt -326 to 440 of genomic DNA*, with an additional 26 bp insertion. |
| *tbb-1(gk207)* | Forward: ggagatgacttcccagaacttg (YJ12636)<br>Reverse: cagaaacacgaagagatggcac (YJ12637)<br>WT: 632 bp<br>MUT:333 bp | Deletion of nt -286 to 13 of genomic DNA*. |
| *lam-1::GFP (qyIs8)* | GFP signal visible under compound microscope | Integrated transgene (Ziel et al., 2009) |

| | | |
|---|---|---|
| *nid-1::mNG (qy38)* | mNG signal visible under compound microscope | mNG insertion at the C-terminus of NID-1 (Keeley et al., 2020) |
| *tac-1(or455)* | Homozygotes viable at 15°C and display embryo lethality at 25°C; maintained at 15°C and phenotype checked at 25°C. | Contains a missense mutation M58I (Bellanger et al., 2007) |

*nt number starting with initiation codon ATG as nt 1-3.

**see Supplemental Table 3 for specifics of edited sequence

**Supplemental Table 3: CRISPR reagents and sequences of edited alleles**

| Allele name | crRNA sequence (bold letters mark **PAM**) (5' to 3') | Genomic sequence flanking the edited site (underlined)* and edited allele sequence** (5' to 3') |
|---|---|---|
| *efa-6(ju1658)* | CGTTTTCAGAGTGATGGCGA**AGG** | CTCGTTTTCAGAGTG/ CGCGACTTTCGCCAT |
| *efa-6(ju1883)* | (a)GAGGCACTGGCCACCATTGA**TGG** (b)TTTGCAACGATCTCCTCGTT**CGG** | TGACGATGACAAGAG[#]/ GAGATCGTTGCAAAA |
| *efa-6(ju1903)* | (a)ACAAGGATGACGATGACAAG**AGA** (b)TTTGCAACGATCTCCTCGTT**CGG** | CAAGGATGACGATGA[#]/ CTTTTTGCAACGATC |
| *efa-6(ju1943)* | (a)TTTGCAACGATCTCCTCGTT**CGG** (b)TCAGAAGTTGCATGGAGTGA**CGG** | ATGAAATCTCCGAAC/ TCCATGCAACTTCTG |
| *efa-6(ju1990)* | (a)TCAGAAGTTGCATGGAGTGA**CGG** (b)GATGCAACTGTGGTACCTGG**AGG** | GACCAACTGCCTCCA/ CTCCATGCAACTTCT |
| *efa-6(ju2026)* | (a)GATGCAACTGTGGTACCTGG**AGG** (b)GTTGATGAAGAACAATTTAT**TGG** | ACTGCCTCCAGGTAC/ AAAAGCTCACCAATA |
| *efa-6(ju1974)* | AAGAGAAATGAGCACATCAC**CGG** | TTTTGTAATGGCTGGTGATGT CCTTATCTCCCTCAATCGGAA CGTTTCTAGCACTTATG |
| *efa-6(ju2011)*/ *efa-6(ju2028)* | AAGAGAAATGAGCACATCAC**CGG** | TTTTGTAATGGATGATGATGT CCTTATCTCCCTCAATCGGAA CGTTTCTAGCACTTATG |
| *efa-6(ju1833)* | (a)GGATCTTCATGGACCAAATA**TGG** (b)GCTGAGTACTCATGAAGTACTCAC**G GG** | GATCTTCATGGACCA/ CGGGTTGCCGAACTA |
| *tba-1(ju1869)* | CATTGAGAGCTCCATCGAAA**CGG** | TCACCGCTTCTCTTTGTTTTG ACGGTGCCCTTAATGTTGAT C |

| | | |
|---|---|---|
| *tba-1(ju1872)* | CATTGAGAGCTCCATCGAAA**CGG** | TCACCGCTTCTCTTGCTTTTGACGGTGCCCTTAATGTTGATC |
| *tba-1(ju1876)* | CATTGAGAGCTCCATCGAAA**CGG** | TCACCGCTTCTCTTAAGTTTGACGGTGCCCTTAATGTTGATC |
| *tba-1(ju1871)* | CATTGAGAGCTCCATCGAAA**CGG** | TCACCGCTTCTCTTGAGTTTGACGGTGCCCTTAATGTTGATC |
| *tba-2(ju1875)* | ACTGCTTCCTTGAGATTCGA**TGG** | CCTCAATCACCGCCTCTTTATGTTTTGACGGTGCCCTCAACG |
| *tba-1(ju1868)* | CATTGAGAGCTCCATCGAAA**CGG** | TCACCGCTTCTCTTTGTTTTGACGGTGCCCTTAATGTTGATC |

* genomic sequences flanking the edited sites are underlined and separated by / for 5' and 3' flanking sequences.

** red letters are edited nucleotides that cause desired amino acid changes; green letters are synonymous changes

[#] 5' flanking sequences are within the GFP::FLAG knock in sequences in *efa-6(ju1658)*.

**References to Supplemental Tables:**

Antoshechkin, I., Han, M., 2002. The *C. elegans evl-20* gene is a homolog of the small GTPase ARL2 and regulates cytoskeleton dynamics during cytokinesis and morphogenesis. Dev Cell 2, 579-591.

Bellanger, J.M., Carter, J.C., Phillips, J.B., Canard, C., Bowerman, B., Gonczy, P., 2007. ZYG-9, TAC-1 and ZYG-8 together ensure correct microtubule function throughout the cell cycle of *C. elegans* embryos. J Cell Sci 120, 2963-2973.

Chuang, C.H., Schlientz, A.J., Yang, J., Bowerman, B., 2020. Microtubule assembly and pole coalescence: early steps in *Caenorhabditis elegans* oocyte meiosis I spindle assembly. Biol Open 9.

Dejima, K., Hori, S., Iwata, S., Suehiro, Y., Yoshina, S., Motohashi, T., Mitani, S., 2018. An Aneuploidy-Free and Structurally Defined Balancer Chromosome Toolkit for *Caenorhabditis elegans*. Cell Rep 22, 232-241.

Dickinson, D.J., Pani, A.M., Heppert, J.K., Higgins, C.D., Goldstein, B., 2015. Streamlined Genome Engineering with a Self-Excising Drug Selection Cassette. Genetics 200, 1035-1049.

Honda, Y., Tsuchiya, K., Sumiyoshi, E., Haruta, N., Sugimoto, A., 2017. Tubulin isotype substitution revealed that isotype combination modulates microtubule dynamics in *C. elegans* embryos. J Cell Sci 130, 1652-1661.

Keeley, D.P., Hastie, E., Jayadev, R., Kelley, L.C., Chi, Q., Payne, S.G., Jeger, J.L., Hoffman, B.D., Sherwood, D.R., 2020. Comprehensive Endogenous Tagging of Basement Membrane Components Reveals Dynamic Movement within the Matrix Scaffolding. Dev Cell 54, 60-74 e67.

Sallee, M.D., Zonka, J.C., Skokan, T.D., Raftrey, B.C., Feldman, J.L., 2018. Tissue-specific degradation of essential centrosome components reveals distinct microtubule populations at microtubule organizing centers. PLoS Biol 16, e2005189.

Waaijers, S., Munoz, J., Berends, C., Ramalho, J.J., Goerdayal, S.S., Low, T.Y., Zoumaro-Djayoon, A.D., Hoffmann, M., Koorman, T., Tas, R.P., Harterink, M., Seelk, S., Kerver, J., Hoogenraad, C.C., Bossinger, O., Tursun, B., van den Heuvel, S., Heck, A.J., Boxem, M., 2016. A tissue-specific protein purification approach in Caenorhabditis elegans identifies novel interaction partners of DLG-1/Discs large. BMC Biol 14, 66.

Wang, S., Tang, N.H., Lara-Gonzalez, P., Zhao, Z., Cheerambathur, D.K., Prevo, B., Chisholm, A.D., Desai, A., Oegema, K., 2017. A toolkit for GFP-mediated tissue-specific protein degradation in C. elegans. Development 144, 2694-2701.

Wang, S., Wu, D., Quintin, S., Green, R.A., Cheerambathur, D.K., Ochoa, S.D., Desai, A., Oegema, K., 2015. NOCA-1 functions with gamma-tubulin and in parallel to Patronin to assemble non-centrosomal microtubule arrays in *C. elegans*. Elife 4, e08649.

Zheng, Q., Schaefer, A.M., Nonet, M.L., 2011. Regulation of *C. elegans* presynaptic differentiation and neurite branching via a novel signaling pathway initiated by SAM-10. Development 138, 87-96.

Ziel, J.W., Hagedorn, E.J., Audhya, A., Sherwood, D.R., 2009. UNC-6 (netrin) orients the invasive membrane of the anchor cell in *C. elegans*. Nat Cell Biol 11, 183-189.
